# Illuminating Dark Chemical Matter using the Cell Painting Assay

**DOI:** 10.1101/2023.05.31.542818

**Authors:** Axel Pahl, Jie Liu, Sohan Patil, Soheila Rezaei Adariani, Beate Schölermann, Jens Warmers, Jana Bonowski, Sandra Koska, Sonja Sievers, Slava Ziegler, Herbert Waldmann

## Abstract

The identification of bioactive small molecules is at the heart of chemical biology and medicinal research. The screening for modulators of disease-relevant targets and phenotypes is the first step on the way to new drugs. Therefore, large compound libraries have been synthesized and employed by academia and, particularly, pharmaceutical companies to meet the need for chemical entities that are as diverse as possible. Extensive screening of these compound libraries revealed a portion of small molecules that is inactive in more than 100 different assays and was therefore termed ‘dark chemical matter’ (DCM). Deorphanization of DCM promises to yield very selective compounds as they, by definition, should have less off-target effects. We employed morphological profiling using the Cell painting assay (CPA) to detect bioactive DCM compounds. CPA is not biased to a given target or phenotype and can detect various unrelated mechanisms and modes of action. Within the DCM collection, we identified bioactive compounds and confirmed several modulators of microtubules, DNA synthesis and pyrimidine biosynthesis. Profiling approaches are therefore powerful tools to probe compound collections for bioactivity in an unbiased manner and particularly suitable for deorphanization of DCM.

## Introduction

Screening of large compound collections using target or phenotype-based assays is usually the initial step in the discovery of biologically active small molecules. Typically, 0.1 to 1 % of these compounds scores as hits in such screens and higher hit rates are undesirable as they are often linked to the detection of promiscuous compounds (frequent hitters), e.g. polypharmacological or pan-assay interference compounds (PAINS).^1, 2^ This raises the question of what are the right small molecules to synthesize with regard to bioactivity. Different organic-synthesis design principles proved powerful for the identification of small-molecule modulators of biological targets, like biology-oriented synthesis (BIOS)^3, 4^, diversity-oriented synthesis (DOS) ^5^ or pseudo-natural products^6, 7^. Generally, however, screening collections are often compiled based on chemical diversity and attractiveness, considering physicochemical and or drug-like properties.^8^ Analysis of the biological profiles of compound collections by Wassermann et al. identified a set of compounds from the Novartis and NIH Molecular Libraries Program screening collections that were consistently inactive in at least 100 biological assays.^8^ This set of compounds was termed ‘dark chemical matter’ (DCM), a term initially coined by A. Pope.^9^ Interestingly, DCM do not comprise separate scaffold classes that have been found inactive in all these assays. In contrast, DCM compounds are often close analogues of biologically active small molecules, have drug-like physicochemical properties, tend to have higher aqueous solubility and are less hydrophobic than active compounds.^8^ Hence, DCM is endowed with drug-like features for modulation of biological targets. Among the possible reasons for the lack of bioactivity may be the focus on a narrow drug-target space, mostly considering mammalian targets and inappropriate screening concentrations.^8^ Conducting an increasing number of focused screening assays could identify some active DCM^10^, however, these assays are biased towards the target or phenotype of interest. Profiling approaches, in contrast, can capture modulation of multiple targets simultaneously and identify perturbed targets in a less biased manner. In fact, Wassermann et al. detected changes in the expression of 61 selected genes by DCM^8^, indicating that, indeed, DCM does not represent completely biologically inert compounds. Therefore, unbiased profiling appears to be particularly suitable for the analysis of DCM. Here we report on the use of morphological profiling by means of the Cell Painting Assay (CPA) for deorphanization of DCM. A set of ca. 7,700 DCM compounds was profiled using CPA and suggested modes of action for various active small molecules. In particular, we identified inhibitors of microtubule dynamics, DNA synthesis or *de novo* pyrimidine synthesis, thus confirming that DCM may not be biologically inert. Hence, morphological profiling indeed can be used for deorphanization of DCM and can identify highly selective compounds as DCM by definition is expected to exhibit less polypharmacology.

## Results and Discussion

We employed CPA to profile a subset of DCM compounds (see below). CPA is a morphological profiling approach^11, 12^, in which cells are exposed to perturbagens prior to staining with different dyes for visualization of different cell components and compartments, i.e., DNA, RNA and nucleoli, endoplasmic reticulum, actin cytoskeleton, Golgi, plasma membrane and mitochondria. High-content imaging and analysis leads to the extraction of hundreds of morphological features using CellProfiler (https://cellprofiler.org/13). The morphological feature profiles are normalized to DMSO controls and feature values are calculated as Z-scores. In our set-up, CPA profiles are composed of 579 features.^14^ The percentage of significantly changed features is defined as *induction* and used as a measure of bioactivity (compounds with induction ≥ 5% are considered to be active). Profiles sharing similarity (also called biosimilarity) of ≥ 75 % are considered similar. By comparing the profiles of active compounds to those of reference compounds with annotated activities, target hypotheses can be derived (details in the “Methods” section). In addition, the similarity to thus far twelve biological clusters can be determined by comparing CP subprofiles.^15^

Using Python and the cheminformatics toolkit RDKit (https://www.rdkit.org), the set of 19,976 compounds of the ChemDiv DCM library was standardized for the largest fragment using the provided SD file and InChIKeys were added. Two InChIKeys could not be generated (leaving 19,974 compounds). In addition to the structure, the original file contained an identifier and the available amount in milligram (”available”). The structural overlap of the full ChemDiv dataset with our internal compound database (“COMAS DB”, 194,832 compounds) was determined by InChIKey to be 2,818 compounds (Figure 1A). The full ChemDiv DCM dataset was filtered for compounds with sufficient availability (”available” ≥ 2 mg), leaving 18,411 compounds. The number of heavy atoms (”NumHA”) and mol weight (”MW”) were calculated and structures with less than 25 NumHA were removed, leaving 10,640 compounds. The structures overlapping with the COMAS DB were removed, leaving 9,254 entries. Using RDKit, a diversity selection based on Morgan fingerprints was performed (Figure S1B) and the first 5,600 compounds were ordered. 5,574 of those were tested together with the available overlapping internal compounds (2,110) in the CPA. In total, 7,684 were subjected to the morphological profiling at a concentration of 10 µM (Figure 1A).

**Figure 1:**
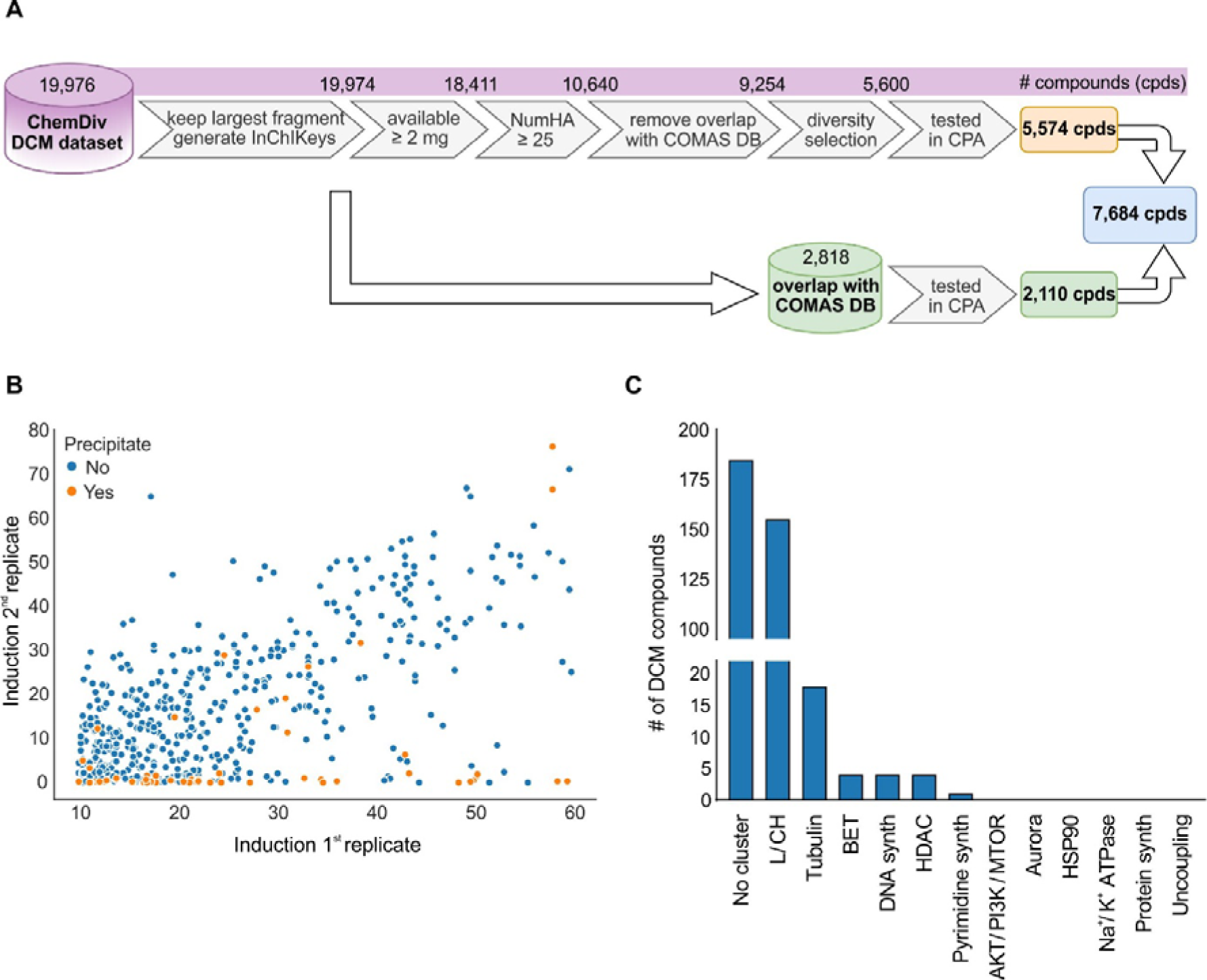
Selection of DCM compounds for the morphological profiling. (A) Summary of the DCM selection process. See also Figure S1. (B) Reproducibility of the induced changes in CPA as determined by the induction value of two replicates for 550 DCM compounds that were initially identified as active in CPA. Correlation between the replicates: r^2^=0.0246. (C) Subprofile analysis. Similarities for the 371 CPA-active DCM compounds to the twelve defined bioactivity clusters. Subprofile biosimilarity ≥ 80% is considered. Synth: synthesis. See also Figure S1 and S2.

Of the 7,684 tested compounds, 942 (12%) showed a significant morphological change compared to the DMSO controls (induction ≥ 5%). 550 compounds with induction ≥ 10% were tested again in CPA. Surprisingly, the activity of only 371 of the initial active compounds was confirmed. This rather low reproducibility is in stark contrast to the CP data with our in-house compounds. When comparing the induction values between two biological replicates, the correlation for our internal compounds was r^2^=0.617, whereas for DCM, r^2^= 0.0246 (see Figure 1B and Figure S2A and S2B). This difference was even more striking when analyzing the profile biosimilarity of the biological replicates: the median biosimilarity was 87% for the internal data set and 78% for DCM. Within our internal CP data, profile biosimilarity of the replicates was ≥ 80% for 78% of the compounds, while only 43% of the DCM set displayed profile similarity of the replicates ≥ 80% (see Figure S2C). By inspection of compound solutions using Tube Auditor we noticed precipitates for a substantial proportion (9.6%) of the initially active DCM compounds that most likely account for the low reproducibility in CPA (Figure 1B and S2A). For comparison, precipitation concerned only 0.4% of our internal compounds. Of note, freshly dissolved DCM compounds apparently did not show precipitates, whereas precipitation occurred after a freeze-thaw cycle. Therefore, the inactivity of at least some of the DCM compounds may be attributed to low DMSO solubility. The CPA profiles of the 371 confirmed compounds were submitted to a cluster subprofile analysis to map similarity to modulation of AKT/PI3K/MTOR, Aurora kinases, BET, DNA synthesis, HDAC, HSP90, lysosomotropism/cholesterol homeostasis (L/CH), Na^+^/K^+^ ATPase, protein synthesis, pyrimidine synthesis, tubulin or uncoupling of the mitochondrial proton gradient.^15^. For this, bioactivity clusters were defined based on profile similarity of, mostly, annotated compounds. Common features of the defining profiles were extracted to construct a median profile for each cluster, i.e., cluster profile which has a reduced number of features as compared to the full profile.^15^ Most of the CPA-active DCM compounds were similar to the lysosomotropism/cholesterol homeostasis cluster (in total 155 small molecules). Moreover, the profiles of several compounds were biosimilar to the clusters of tubulin, DNA synthesis, pyrimidine synthesis or BET modulators (Figure 1C, see also Table S1). The L/CH cluster compiles small molecules that impair cholesterol homeostasis by either specific target modulation or due to their lysosomotropic activity.^16^ Lysosomotropic compounds are weakly basic, lipophilic molecules that share similar physicochemical properties, i.e., logP > 2 and pKa between 6.5 and 11.^17^ Indeed, 67 % of DCM that is similar to L/CH shares properties of lysosomotropic compounds (Figure S3).

Several DCM compounds displayed similarity to the tubulin cluster (Table 1). To confirm the target hypothesis, the influence of the compounds on mitosis was explored as impairment of microtubule dynamics is linked to mitotic arrest. Indeed, the compounds increased the percentage of metaphase cells which were detected using phospho-histone H3 as a marker (Figure 2A). Moreover most compounds almost completely suppressed *in vitro* tubulin polymerization (Figure 2B) and disturbed the microtubule cytoskeleton in cells (Figure 2C and Figure S4). Of note, target prediction based on chemical similarity using the Similarity Ensemble Approach (SEA, https://sea.bkslab.org)^18^ revealed tubulin as a potential target for compounds **1, 2, 3** and **4**. However, for compounds **1, 3** and **4**, tubulin was not among the top 20 predictions. Only for compound **2** tubulin was ranked among the top 5 predicted targets and would have been considered for validation experiments. The tubulin-targeting chemical space is very broad and often only minor structural modifications turn a compound into tubulin modulator^19^ which hampers target prediction based on chemical similarity. These results demonstrate that prediction based on similar morphological profiles and particularly using cluster biosimilarity reliably identifies microtubule targeting agents in compound collections.

**Figure 2.**
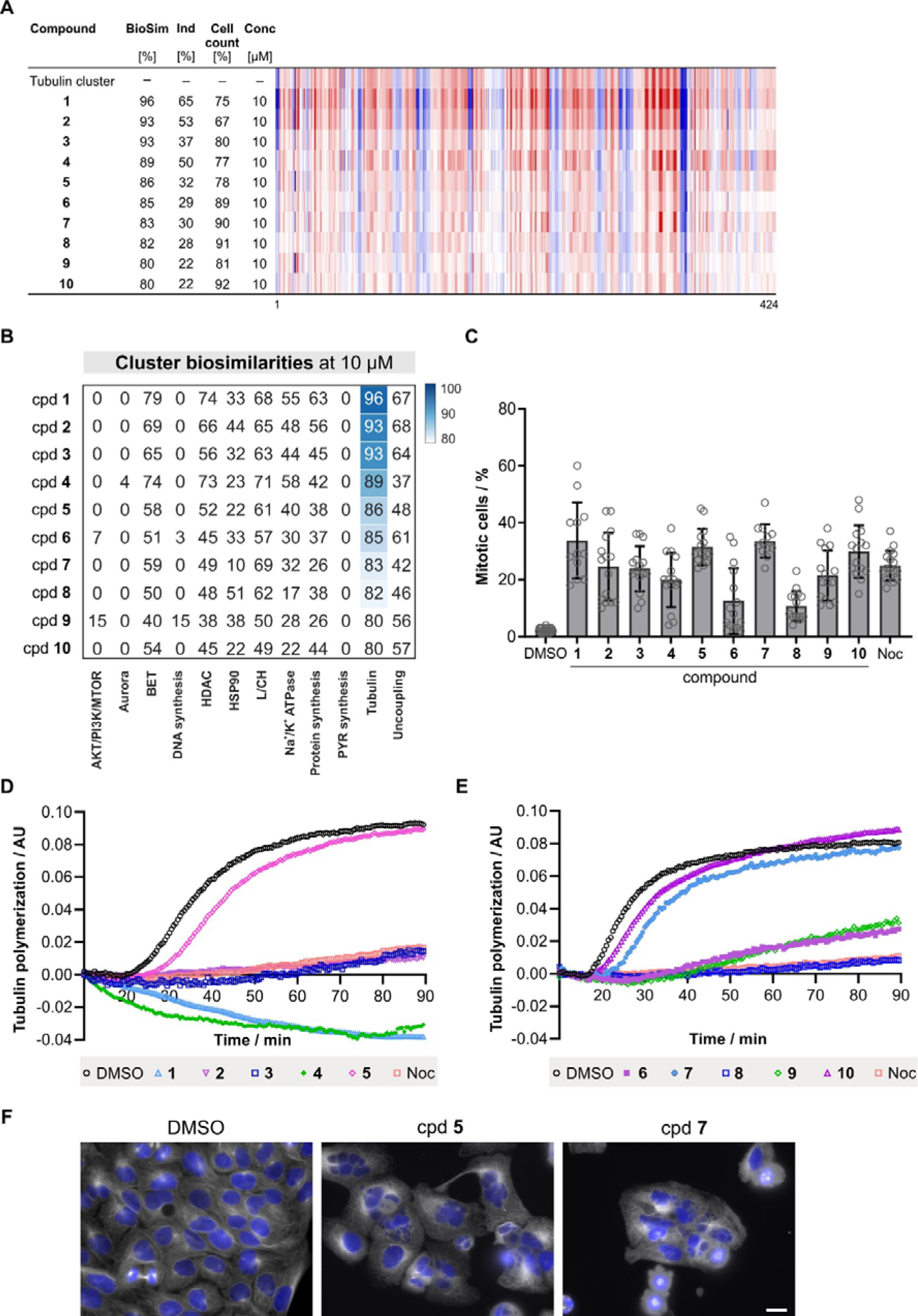
Identification of DCM targeting tubulin. (A) Similarity of compounds (cpds) **1-10** to the tubulin cluster profile. The top line profile is set as a reference profile (100 % biological similarity, BioSim) to which the following profiles are compared. Blue color: decreased feature, red color: increased feature. BioSim: biosimilarity, Ind: induction, Conc: concentration. (B) Cluster biosimilarity heatmap for compounds **1-10**. PYR: pyrimidine. (C) Influence of compounds **1-10** on the number of mitotic cells. U2OS cells were treated with the compounds (30 µM) or 0.1 µM Nocodazole (Noc) for 24 h prior to detection of mitotic cells using anti-phospho-histone H3. Data are mean values ± SD (n = 3). (D and E) Influence of the compounds on the in vitro tubulin polymerization at 20 µM. Nocodazole was used as a control (2 µM). (F) Influence on microtubules in cells. U2OS cells were treated with 30 µM of cpd **5** or cpd **7** or DMSO as a controls for 24 h prior to staining with anti-tubulin antibody (white) or DAPI (blue) to visualize the DNA. Scale bar: 20 µm. See also Figure S4.

**Table 1.**
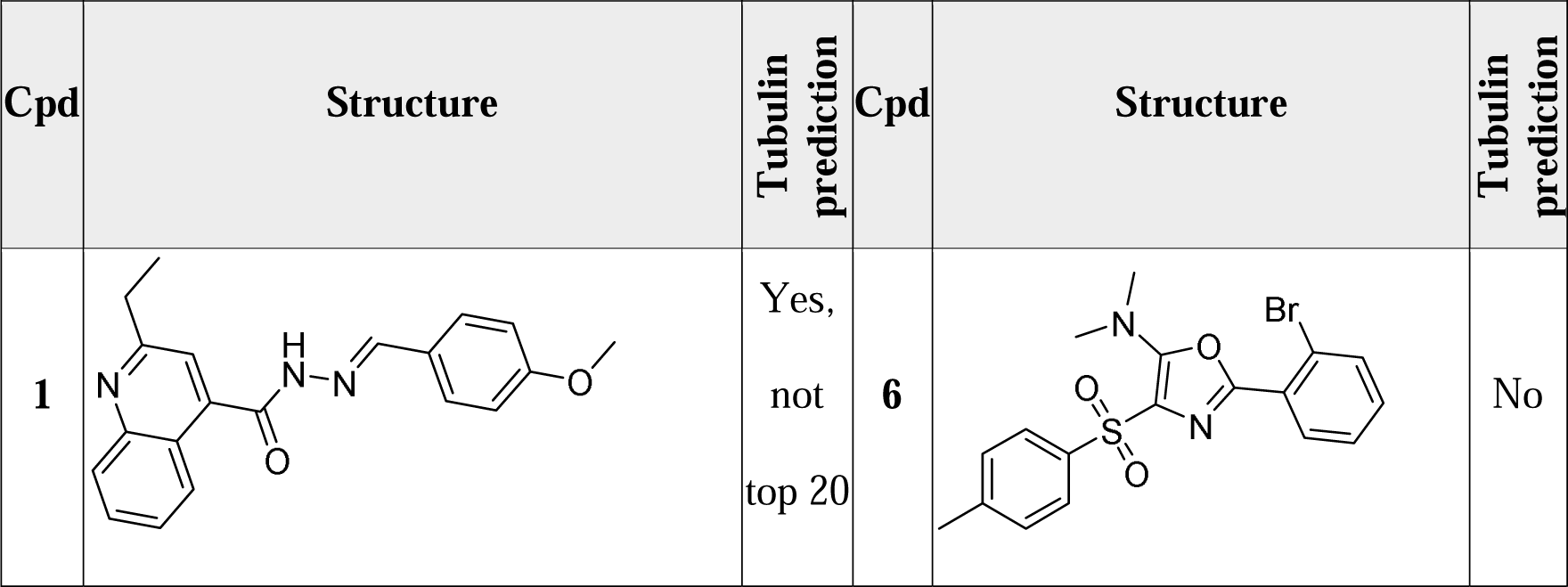
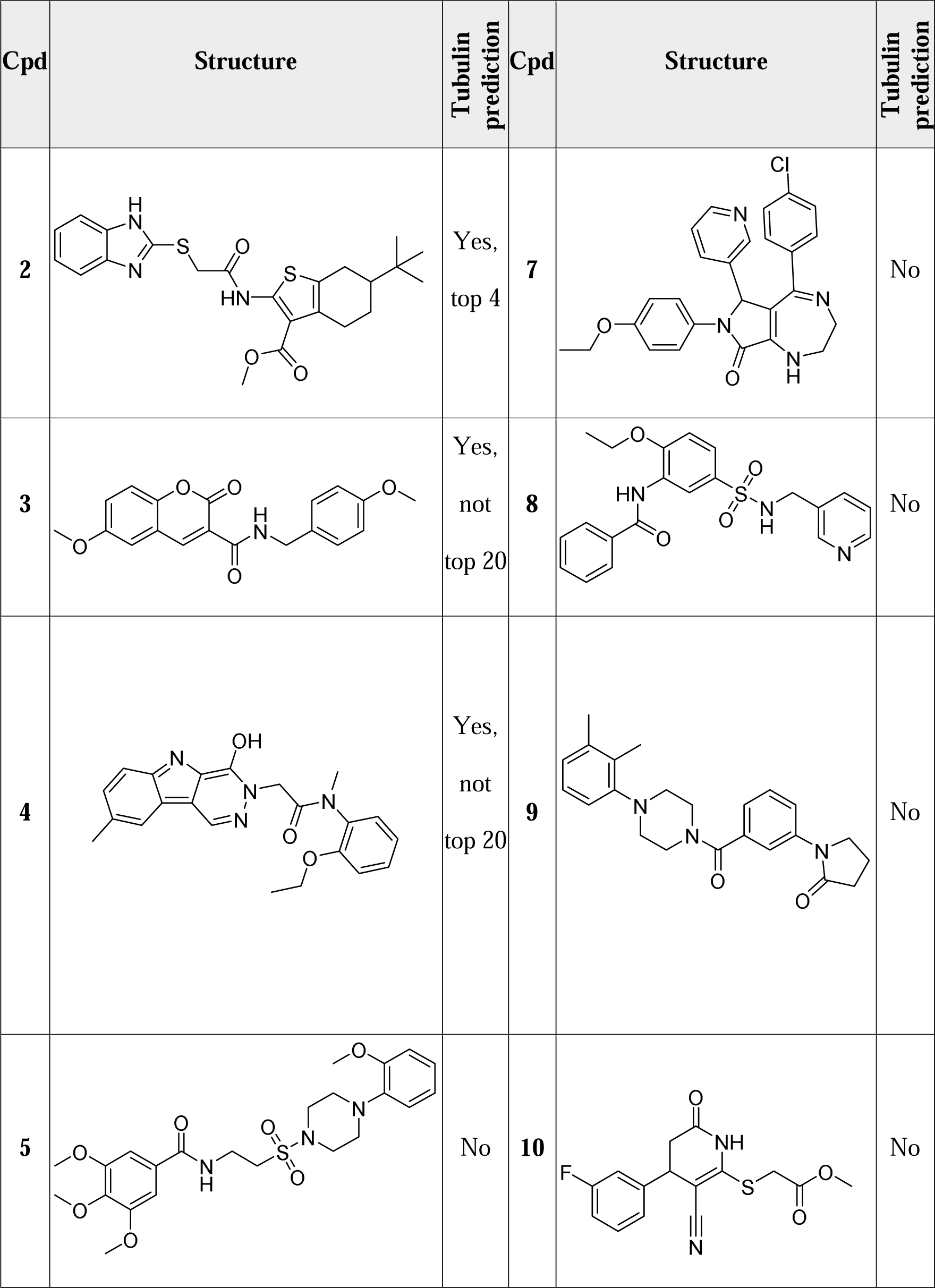
Prediction of tubulin as a target for DCM compounds (cpds) 1-10. The tool SEA (Similarity Ensemble Approach)^18^ was used to detect tubulin as a potential target.

| Cpd | Structure | Tubulin prediction | Cpd | Structure | Tubulin prediction |
| --- | --- | --- | --- | --- | --- |
| **1** |  | Yes,<br>not<br>top 20 | **6** |  | No |
| 2 |  | Yes, top 4 | 7 |  | No |
| 3 |  | Yes, not top 20 | 8 |  | No |
| 4 |  | Yes, not top 20 | 9 |  | No |
| 5 |  | No | 10 |  | No |

We noticed a profile similarity between compound **11** and the microtubule destabilizer Colchicine (Figure 3A and 3B). Although affecting microtubule dynamics, the CPA profile of Colchicine (100 nM) is not similar to the prototypic tubulin inhibitors Nocodazole, Vinblastine or Vincristine (Figure 3C). This is in line with the low similarity of Colchicine to the tubulin cluster profile (61 %, Figure 3B). Instead, Colchicine displays biosimilarity to Epothilone B, which is a microtubule stabilizer^20^, and to Parbendazole, which suppresses tubulin polymerization^21^ (Figure S5A). Compound **11** increased the number of mitotic cells and led to almost complete depolymerization of microtubules in cells (Figure 3D and 3F). A similar phenotype was observed for 100 nM Colchicine (Figure 3F). These findings reveal that different expressions of impaired microtubule dynamics can be captured by CPA that most likely depend on the severity of microtubule disturbance. Increasing the compound concentrations for the explored microtubule-targeting agents reduces the cell count below the threshold of 50% (usually, only profiles with cell count > 50% are analyzed). However, when CPA profiles at higher concentration with a cell count < 50% were considered, profile biosimilarity between Colchicine and Nocodazole, Vinblastine and Vincristine was observed (Figure S5B) and biosimilarity to the tubulin cluster became evident for Colchicine (Figure S5C).

**Figure 3:**
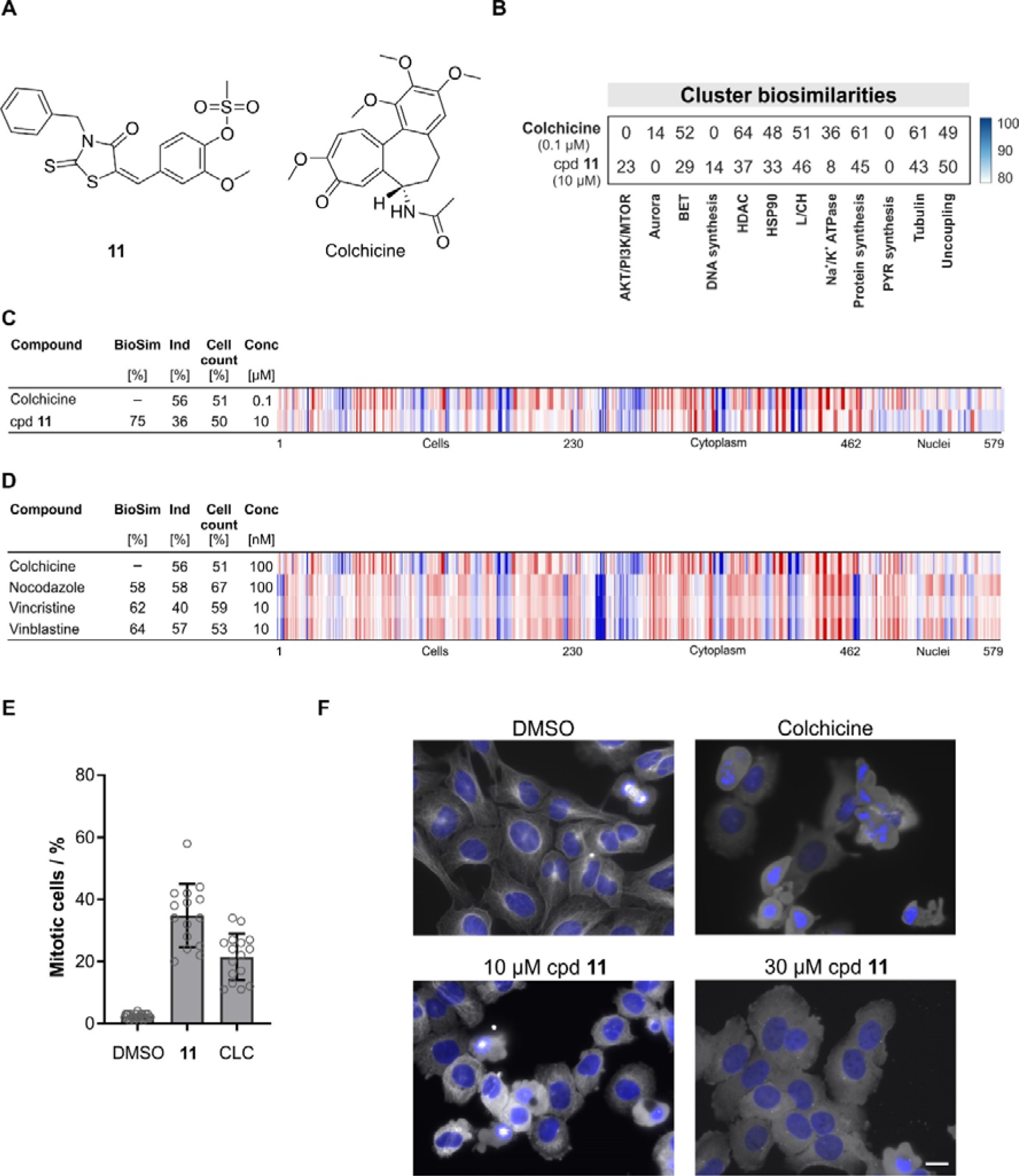
Compound 11 is a microtubule depolymerizer. (A) Structures of compound (cpd) **11** and Colchicine. (B) Cluster biosimilarity heatmap for Colchicine and compound **11**. PYR: pyrimidine. (C) Profile similarity of compound **11** to Colchicine. (D) Profile similarity of Colchicine to Nocodazole, Vincristine and Vinblastine. (C and D) The top line profile is set as a reference profile (100 % biological similarity, BioSim) to which the following profiles are compared. Blue color: decreased feature, red color: increased feature. BioSim: biosimilarity, Ind: induction, Conc: concentration. (E) Influence of compound **11** on the number of mitotic cells. U2OS cells were treated with **11** (10 µM) for 24 h prior to detection of mitotic cells using anti-phospho-histone H3. DMSO and Colchicine (CLC, 0.01 µM) were used as controls. Data are mean values ± SD (n = 3). (F) Influence on microtubules in cells. U2OS cells were treated with 10 or 30 µM of **11** or DMSO and Colchicine (0.01 µM) as controls for 24 h prior to staining with anti-tubulin antibody (white) or DAPI (blue) to visualize the DNA. Scale bar: 20 µm. See also Figure S5.

The nucleoside Vidarabine and three DCM compounds displayed similarity to the DNA synthesis cluster (compound **12-15**, Figure 4A and 4B and Figure S6A). Vidarabine (AraA, **12**, Figure 4A) is an anti-viral agent isolated from *Streptomyces antibioticus* and is known to inhibit viral DNA polymerases.^22^ According to Wasserman *et al.*, Vidarabine is inactive in more than hundred biochemical assays ^8^. In contrast, its epimer adenosine (Figure S6B) is a bioactive compound that plays a role in various processes, e.g., inflammation, vasodilation and angiogenesis.^23^ We recently reported bioactivity for Vidarabine in CPA and profile similarity to the iron chelator Deferoxamine that originates from inhibition of DNA synthesis.^24^ At 10 and 30 µM Vidarabine, induction of 13% and 27 %, respectively, was detected in CPA with high biosimilarity to the DNA synthesis cluster at 30 µM (Figure 4B and Figure S6A). Hence, CPA points towards inhibition of DNA synthesis by Vidarabine in mammalian cells that was experimentally confirmed (Figure 4C). Of note, Adenosine is inactive in CPA up to a concentration of 30 µM (induction < 1%).

**Figure 4:**
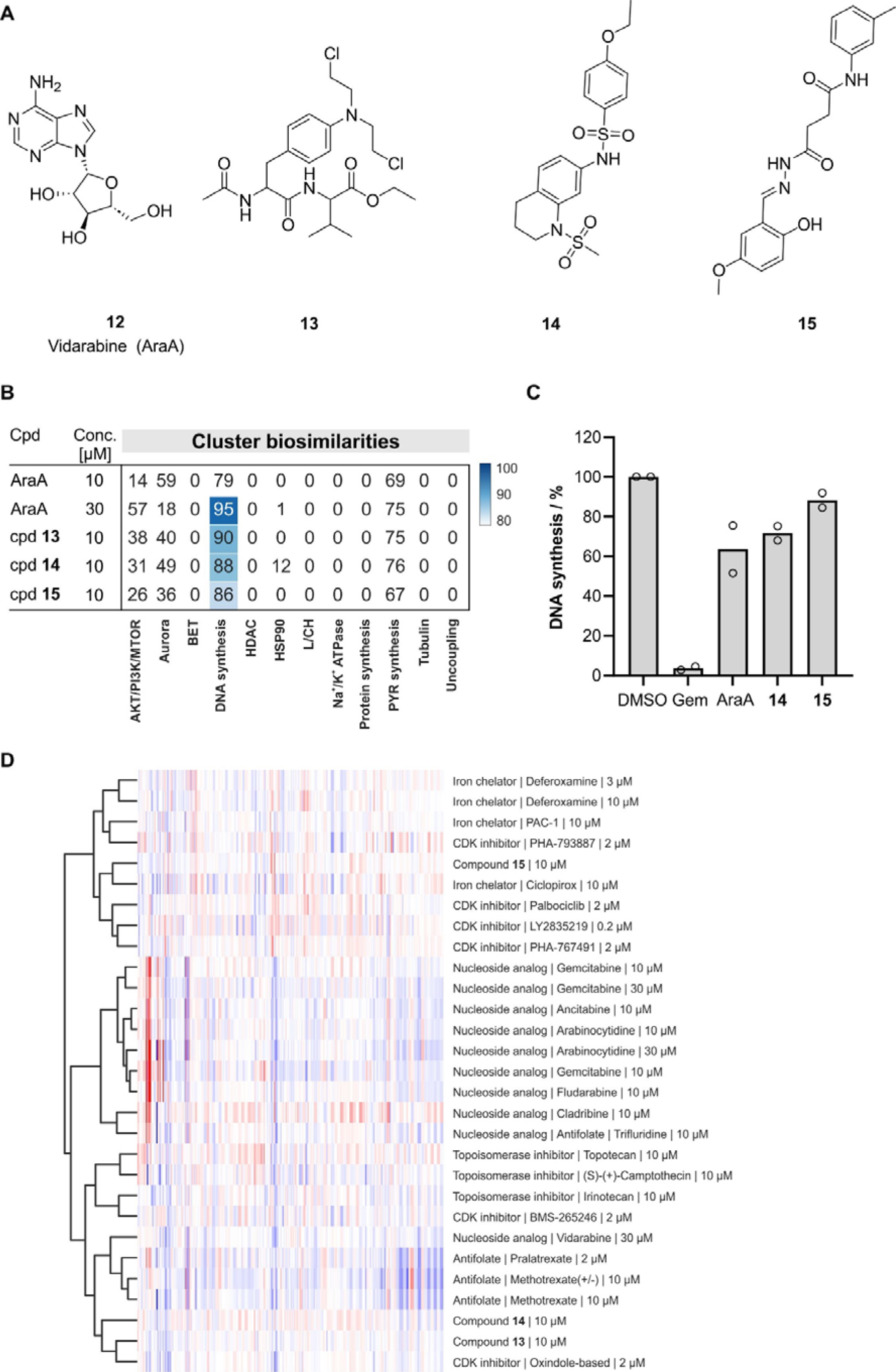
DCM compounds impair DNA synthesis. (A) Structure of DCM compounds **12**- **15** that display biosimilarity to the DNA synthesis cluster. (B) Cluster biosimilarity for Vidarabine (AraA, **12**) and compounds **13-15**. PYR: pyrimidine. (C) Influence on DNA synthesis after treatment of U2OS cells for 1 h followed by EdU pulse for 1h. Gemcitabine (Gem) and DMSO were used as controls. Data are mean values (n=2). (D) Hierarchical clustering using the non-DNA cluster features for compounds of the DNA synthesis cluster and compound **13-15**. Only the 291 non-cluster features were used. See Figure S6F for the structure of the oxindole-based CDK inhibitor^25^. See also Figure S6.

Compound **13** has structural similarity to the alkylating agent Melphalan (Figure S6C) and both compounds share very high biosimilarity (93 %, Figure S6D). Melphalan alkylates purines and blocks DNA synthesis, which is reflected in the high similarity to the DNA synthesis cluster (Figure S6E). A similar mechanism of action is expected for compound **13**. Compound **14** and to a lesser extent compound **15** also inhibited DNA synthesis after 1 h incubation (Figure 4C). Hierarchical clustering using the non-DNA cluster features grouped compound **15** with iron chelators whereas **13** and **14** were located together with nucleosides, antifolates and topoisomerase inhibitors (Figure 4D).

CPA identified the DCM compound **16** as a potential inhibitor of *de novo* pyrimidine biosynthesis as it displayed 84 % similarity to the pyrimidine synthesis cluster (Figure 5A- 5C). As recently reported, this cluster contains modulators of enzymes in pyrimidine biosynthesis like dihydroorotate dehydrogenase (DHODH) and UMP synthase (UMPS) as well as inhibitors of mitochondrial complex III.^26^ **16** suppressed the growth of HCT-116 cells that rely on *de novo* pyrimidine biosynthesis. This influence was rescued in the presence of uridine, which feeds into the salvage pathway and makes *de novo* pyrimidine synthesis dispensable (Figure 5D und 5E). Moreover, **16** dose-dependently inhibited the DHODH activity *in vitro* with an approximate IC_50_ of 7.5 µM (Figure 5F), thus confirming the predicted target.

**Figure 5.**
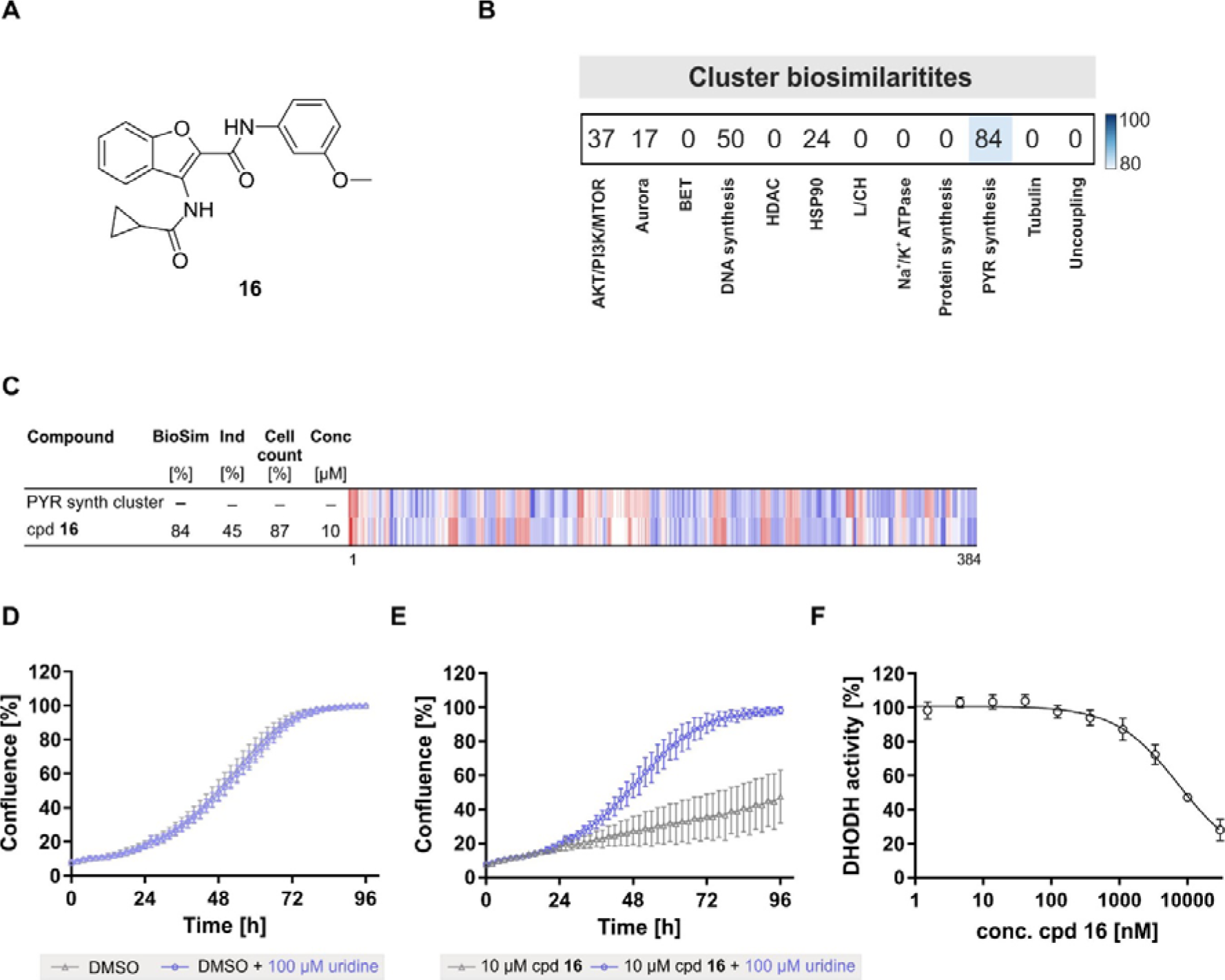
Compound 16 is a DHODH inhibitor. (A) Structure of compound **16**. (B) Similarity of compound (cpd) **16** to the pyrimidine (PYR) synthesis cluster profile. The top line profile is set as a reference profile (100 % biological similarity, BioSim) to which the following profiles are compared. Blue color: decreased feature, red color: increased feature. BioSim: biosimilarity, Ind: induction, Conc: concentration. (C) Cluster biosimilarity for compound **16** (10 µM). PYR: pyrimidine. (D and E) Influence on the growth of HCT-116 cells in absence and presence of 100 µM uridine for DMSO (C) and compound (cpd) **16** (D). Cell confluence was used as a measure of cell growth. Data mean values ± SD (n=3). (F) Influence of **16** on the enzymatic activity of DHODH. Data mean values ± SD (n=3).

Of the 371 DCM compounds that were reproducibly active in CPA, 185 showed no significant similarity to any of the defined clusters. The biological activity of these small molecules can be further assessed in different target- or cell-based assays. As DCM per definition is comprised of compounds that are inactive in at least 100 assays, the choice of the proper setup for assay #101 should be carefully made. In particular, phenotypic assays allow detection of target modulation in cells. Choosing the right system (primary cells), exogenous stimulus (if appropriate) and a suitable biomarker as a readout (endogenous marker) are essential for the physiological relevance of the assay (and are known as the phenotypic screening ‘rule of 3’).^27^ To gain further insight into the MoA for these 185 CP-active DCM compounds, we selected a Hedgehog-dependent osteogenesis differentiation assay using the murine mesenchymal stem cells C3H10T1/2 which employs stimulation with the Hedgehog agonist Purmorphamine^28^, and that complies with this ‘rule of 3’. Osteoblast differentiation was monitored using the activity of the endogenous marker alkaline phosphatase. 24 compounds reduced the activity of alkaline phosphatase without being toxic to the cells with IC_50_ values ranging from 0.5 µM to 7.5 µM (see Figure 6A and Table S2). To evaluate a direct effect on the Hedgehog (Hh) pathway, we employed a GLI-dependent reporter gene assay using the Shh-LIGHT2 cells^29^. Seven compounds (compound **17, 18, 20-24**) dose-dependently reduced GLI-mediated gene expression, hence, they directly influence Hh signaling. Most small molecules that modulate Hh signaling target the seven-pass transmembrane protein Smoothened (SMO)^30^ and target prediction using the SEA tool (https://sea.bkslab.org/)^18^ suggested SMO as a target for compound **17**.

**Figure 6.**
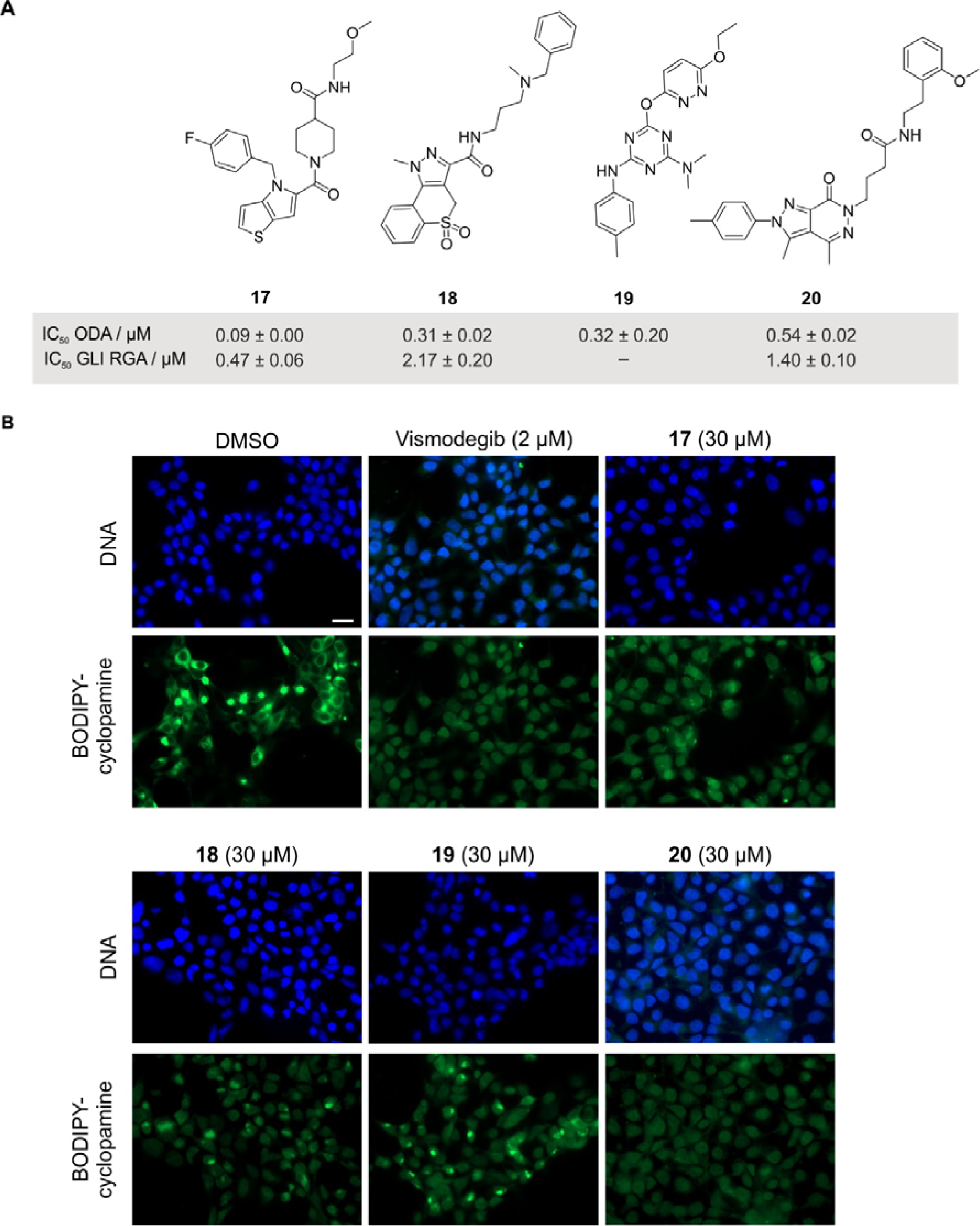
DCM compounds inhibit Hedgehog-dependent osteogenesis. (A) Structure of compounds **17-20** and the corresponding IC_50_ values for inhibition of Purmorphamine- induced osteoblast differentiation (ODA) and GLI-responsive reporter gene assay (GLI RGA). Data are mean values ± SD (n=3). (B) Competition with BODIPY-cyclopamine for binding to SMO. HEK293T cell expressing SMO were fixed and treated with BODIPY-cyclopamine (green) and the compounds for 4 h. The SMO antagonist Vismodegib was used as a control. DAPI was used to visualize the DNA. Images are representative of three biological replicates (n = 3). Scale bar: 20 µm. See also Table S5.

We used the SMO antagonist Cyclopamine labelled with BODIPY to assess potential binding of DCM compounds to SMO. Similar to the SMO antagonist Vismodegib, compounds **17, 18**, and **20** prevented BODIPY-cyclopamine from binding to SMO, indicating that they modulate Hh signaling by antagonizing SMO (Figure 7B). These results are in line with the observed inhibition of GLI-dependent transcription. In contrast, compound **19** did not effectively compete with BODIPY-cyclopamine. This finding, together with the lack of influence on GLI-dependent reporter gene expression, suggests that **19** inhibits Purmorphamine-mediated osteogenesis but may not directly target canonical Hh signaling.

### Discussion and Conclusions

Our results demonstrate that phenotypic screening and profiling approaches are an efficient strategy to uncover biologically active DCM. Subjecting DCM to cellular assays rather than target-based approaches exposes DCM compounds to multiple targets that go beyond protein targets. Moreover, the unbiased nature of the employed morphological profiling identified different bioactivities for various DCM. Testing DCM in an increasing number of assays can identify bioactive DCM as demonstrated by Wassermann et al.^10^ and emphasize that, most likely, these DCM compounds have not been exposed yet to the ‘right’ assay. Surprisingly, several CPA-active DCM compounds target microtubules, which are one of the most frequently addressed (off-) targets of small molecules^19^. Most likely screening activities did not focus on tubulin in the past years and this mechanism of action has remained undetected for DCM. We identified several DCM compounds that interfere with DNA synthesis. Inhibition of DNA replication can be achieved by various mechanisms that target proteins, DNA directly or through chelation of Fe(II)/Fe(III) and CPA is particularly suitable for the detection of this mode of action.^24^. This also applies to the modulation of *de novo* pyrimidine biosynthesis: CPA reliably detects DHODH inhibitors and many of them show only moderate potency in an *in vitro* DHODH activity assay^31^ and may remain below the activity threshold limit of such assay. These examples emphasize the power of morphological profiling to deorphanize DCM as bioactivity is detected in an unbiased manner, which was also demonstrated for gene expression profiling^8, 10^. Employing physiologically-relevant phenotypic assays for screening will further support the bioactivity annotation of DCM as exemplified by the Hedgehog-induced osteogenesis assay, which was employed here. Therefore, DCM should not be regarded as inert chemical entities and should be included in screening campaigns as they are expected to be more selective and to show less polypharmacology^8^.

In summary, we employed morphological profiling by means of CPA to explore the apparently biologically inert DCM. Cell painting proved to be particularly suitable for deorphanization of DCM and identified modulators of microtubule dynamics, DNA synthesis and DHODH. Phenotypic screening of CPA-active compounds in a physiologically relevant osteoblast differentiation assay additionally identified inhibitors of Hedgehog signaling that target Smoothened. Therefore, exploration of DCM collections in less target-biased phenotypic assays or unbiased profiling approaches promises to detect biologically active DCM compounds, which are expected to exhibit less polypharmacology and may deliver highly selective compounds for chemical biology and medicinal chemistry research.

## EXPERIMENTAL SECTION

### Materials

Shh-LIGHT2 cells ^29^ were maintained in high-glucose DMEM, 10% heat inactivated fetal calf serum, 1 mM sodium pyruvate, 3 mM L-glutamine, 5 mM Hepes pH 7.4, 100 U/mL penicillin and 0.1 mg/mL streptomycin. The human kidney cell line HEK293T (ATCC, CRL-11268; RRID:CVCL_1926), human osteosarcoma U2OS cells (CLS #300364; RRID:CVCL_0042) and human colorectal cancer cell line HCT-116 (DSMZ #581; RRID:CVCL_0291) were cultured in Dulbecco’s Modified Eagle’s medium (DMEM with 4.5 g/L Glucose, L-glutamine and 3.7 g/L sodium bicarbonate; PAN Biotech, #P04-03550) supplemented with 10% of fetal bovine serum (FBS, Invitrogen, cat# 10500-084), 1 mM sodium pyruvate (PAN, #P04-43100) and 1% MEM-non-essential amino acids (PAN, #P08-32100). The cell lines were cultured in a humidified atmosphere at 37°C and 5% CO_2_. All cells were regularly checked for mycoplasma contaminations and cells were found to be free of contaminations at all times.

### Cell Painting Assay

The Cell Painting Assays follows closely the method described by Bray et al.^32^ as recently reported^15^. “Initially, 5 µL U2OS medium were added to each well of a 384-well plate (PerkinElmer CellCarrier-384 Ultra). Subsequently, U2OS cell were seeded with a density of 1600 cells per well in 20 µL medium. The plate was incubated for 10 min at the ambient temperature, followed by an additional 4 h incubation (37 °C, 5% CO_2_). Compound treatment was performed with the Echo 520 acoustic dispenser (Labcyte). Different concentrations of DMSO were used as controls dependent on the used compound concentration, e.g., 0.1 % DMSO was used as a control for the profiling of compounds at 10 µM. Samples at a given compound concentration were compared to the DMSO sample of the same DMSO concentration. Incubation with compound was performed for 20 h (37 °C, 5% CO_2_). Subsequently, mitochondria were stained with Mito Tracker Deep Red (Thermo Fisher Scientific, Cat. No. M22426). The Mito Tracker Deep Red stock solution (1 mM) was diluted to a final concentration of 100 nM in prewarmed medium. The medium was removed from the plate leaving 10 µL residual volume and 25 µL of the Mito Tracker solution were added to each well. The plate was incubated for 30 min in darkness (37 °C, 5% CO_2_). To fix the cells 7 µL of 18.5 % formaldehyde in PBS were added, resulting in a final formaldehyde concentration of 3.7 %. Subsequently, the plate was incubated for another 20 min in darkness (RT) and washed three times with 70 µL of PBS. (Biotek Washer Elx405). Cells were permeabilized by addition of 25 µL 0.1% Triton X-100 to each well, followed by 15 min incubation (RT) in darkness. The cells were washed three times with PBS leaving a final volume of 10 µL. To each well 25 µL of a staining solution were added, which contains 1% BSA, 5 µL/mL Phalloidin (Alexa594 conjugate, Thermo Fisher Scientific, A12381), 25 µg/mL Concanavalin A (Alexa488 conjugate, Thermo Fisher Scientific, Cat. No. C11252), 5 µg/mL Hoechst 33342 (Sigma, Cat. No. B2261-25mg), 1.5 µg/mL WGA-Alexa594 conjugate (Thermo Fisher Scientific, Cat. No. W11262) and 1.5 µM SYTO 14 solution (Thermo Fisher Scientific, Cat. No. S7576). The plate is incubated for 30 min (RT) in darkness and washed three times with 70 µL PBS. After the final washing step, the PBS was not aspirated. The plates were sealed and centrifuged for 1 min at 500 rpm.

The plates were prepared in triplicates with shifted layouts to reduce plate effects and imaged using a Micro XL High-Content Screening System (Molecular Devices) in 5 channels (DAPI: Ex350-400/ Em410-480; FITC: Ex470-500/ Em510-540; Spectrum Gold: Ex520-545/ Em560-585; TxRed: Ex535-585/ Em600-650; Cy5: Ex605-650/ Em670-715) with 9 sites per well and 20x magnification (binning 2).

The generated images were processed with the *CellProfiler* package (https://cellprofiler.org/, version 3.0.0)^13^ on a computing cluster of the Max Planck Society to extract 1716 cell features per microscope site. The data was then further aggregated as medians per well (9 sites -> 1 well), then over the three replicates.

Further analysis was performed with custom *Python* (https://www.python.org/) scripts using the *Pandas* (https://pandas.pydata.org/) and *Dask* (https://dask.org/) data processing libraries as well as the *Scientific Python* (https://scipy.org/) package.

From the total set of 1716 features, a subset of highly reproducible and robust features was determined using the procedure described by Woehrmann et al.^33^ in the following way: Two biological repeats of one plate containing reference compounds were analysed. For every feature, its full profile over each whole plate was calculated. If the profiles from the two repeats showed a similarity >= 0.8 (see below), the feature was added to the set.

This procedure was only performed once and resulted in a set of 579 robust features out of the total of 1716 that was used for all further analyses.

The phenotypic profiles were compiled from the Z-scores of all individual cellular features, where the Z-score is a measure of how far away a data point is from a median value.

Specifically, Z-scores of test compounds were calculated relative to the Median of DMSO controls. Thus, the Z-score of a test compound defines how many MADs (Median Absolute Deviations) the measured value is away from the Median of the controls as illustrated by the following formula:

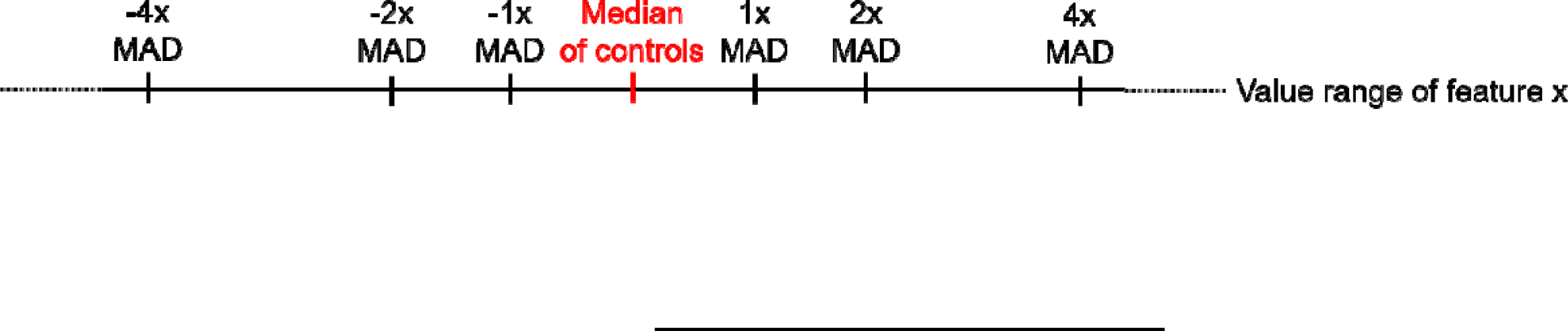

The phenotypic compound profile is then determined as the list of Z-scores of all features for one compound.

In addition to the phenotypic profile, an induction value was determined for each compound as the fraction of significantly changed features, in percent:

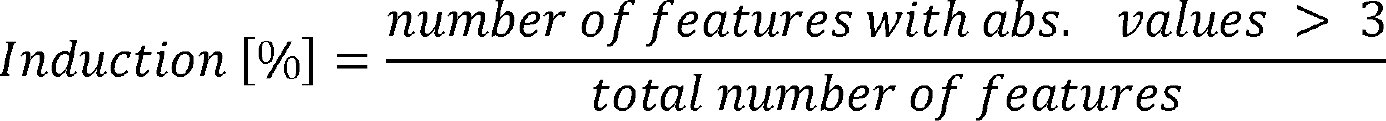

Similarities of phenotypic profiles (termed *Biosimilarity*) were calculated from the correlation distances (CD) between two profiles (https://docs.scipy.org/doc/scipy/reference/generated/scipy.spatial.distance.correlation.html)

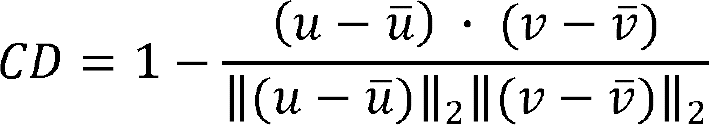

where 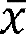 is the mean of the elements of *x, x· y* is the dot product of *x* and *y,* and ||*x*||_2_ is the Euclidean norm of *x*:

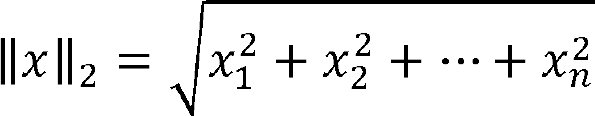

The Biosimilarity is then defined as:

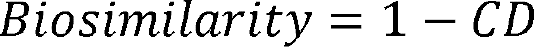

Biosimilarity values smaller than 0 are set to 0 and the Biosimilarity is expressed in percent (0-100).”

### Immunocytochemistry

5,000 U2OS cells were seeded per well in a 96 well plates (Cellvis P96-1-N, 0.13-0.16 mm thickness) and incubated overnight (37 °C, 5% CO_2_). Cells were treated with compounds or DMSO as a control for 24 h. Cells were then fixed using 3.7% paraformaldehyde in PBS and permeabilized with 0.1% Triton X100 (in PBS), and then washed with PBS-T. Subsequently, 100 μL 2% BSA in PBS-T was added to the cells and incubated for 1 h prior to staining with DAPI (Sigma Aldrich, D9542-10MG, 1:1000 dilution) to visualize DNA and anti-tubulin-FITC antibody (Thermo Fisher, MA119581, 1:500 dilution) or anti-phospho-histone H3 antibody (Cell Signaling, #8481, 1:500 dilution) overnight at 4 °C. Images were acquired using Observer Z1 (Carl Zeiss, Germany) using 40X objectives (LD Plan-Neofluar). Moreover, Axiovert 200M microscope (Carl Zeiss, Germany) equipped with 10X objective was used to detect and quantify phospho-histone H3-positive cells using the software MetaMorph 7. For the automated image analysis, the percentage of phospho-histone H3- positive cells was determined using the DNA stain to assess the total number of cells by the software CellProfiler. Data presented are mean values (N = 3) ± SD and are representative of three independent experiments.

### *In vitro* tubulin polymerization assay

In vitro tubulin polymerization assay was performed as described previously Akbarzadeh et al.^35^. Porcine α/β-tubulin (Cytoskeleton, T240-B) was diluted in a general buffer containing 80 mM PIPES (pH 6.9), 2 mM MgCl_2_ and 0.5 mM EGTA. Next, α/β-tubulin (final concentration 10 µM) was added to a solution containing MgCl_2_ and glutamate (Sigma Aldrich, 49621-250G) with a final concentration of 0.88 µM and 0.8 mM, respectively, on a 96-well plate (Corning 3696). Subsequently, compounds at a final concentration of 20 µM were added to the tubulin solution and incubated at room temperature for 20 min. The plate was then incubated on ice for another 20 min followed by the addition of GTP (Thermo Fisher, R0461) to a final concentration of 500 µM. Tubulin polymerization was monitored for 60 min by means of turbidity measurements at 340 nm using Infinite M200 plate reader (Tecan). Data shown is representative of three independent experiments.

### DNA synthesis assay

DNA synthesis was assessed using Click-iT EdU Proliferation Assay for Microplates (C10499, Thermo Fisher Scientific Inc., USA) according to the manufacturer’s instructions. 5,000 U2OS cells were seeded per well into a black 96-well plate and incubated overnight at 37 °C and 5% CO_2_. The next day, the cells were treated with the compounds or DMSO as a control for 1 h. To directly quantify DNA synthesis, the nucleoside analog EdU (5-ethynyl-2’- deoxyuridine, 10 µM) was added to live cells and incubated for an additional 1 h. Incorporation of EdU into newly synthesized DNA was detected upon click reaction and coupling to horseradish peroxidase (HRP). The Amplex UltraRed Reagent was added and its conversion by HRP into a highly fluorescent product was recorded at ex/em 550/595 nm using a Spark plate reader (Tecan). To quantify the total amount of DNA relative to the number of cells, the cells’ nuclei were stained with DAPI (Sigma Aldrich, #10236276001) in a phosphate-buffered saline (PBS) overnight at 4 °C. The following day, detection and quantification of DAPI positive cells was achieved by using an Axiovert 200M microscope (Carl Zeiss, Germany) equipped with 10x objective and the images were processed using the software MetaMorph 7.

### DHODH enzymatic assay

The N-truncated form of hDHODH (aa31−395) was expressed and purified as recently reported Scholermann et al.^31^. The *in vitro* DHODH assay was performed as according to Scholermann et al.^31^. Briefly, the assay monitored the reduction of 2,6- dichlorophenolinindophenol (DCPIP) that was coupled to the oxidation of dihydroorotate by DHODH. 40 µL of purified DHODH (aa31−395, final concentration: 1.25 µg/mL) in assay buffer (50 mM Tris pH 8,0, 150 mM KCl, 0,1% Triton X-100) was incubated with 10 µL of the compounds for 30 min at 37°C followed by 15 min at room temperature. The reaction was initiated by adding 2 mM L-dihydroorotic acid, 0.2 mM decylubiquonone, 0.12 mM 2,6 dichlorphenolindophenol (DCPIP) in assay buffer. DCPIP reduction was monitored at 600 nm using Tecan Spark plate reader for 30 min. The slope of the linear curves over 5 min was calculated to determine the inhibitory activity and IC_50_ values. Values were normalized to the DMSO control and the data were analysed using GraphPad Prism 9 software.

### Cell growth determination and uridine rescue

2,000 HCT-116 cells were seeded per well in a clear 96-well plate and incubated overnight. The next day, the medium was replaced by medium containing the compound or DMSO in presence or absence of 100 µM uridine. Cell growth was monitored by means real-time live-cell analysis with IncuCyte S3 (Essen BioScience). Images were acquired every 2 h for a period of 96 h after treatment. The cell confluence was quantified as a measure of cell growth using the IncuCyte S3 software (Essen BioScience).

### Osteoblast differentiation assay

For assaying signal transduction through the Hh pathway, mouse embryonic mesoderm fibroblast C3H10T1/2 cells were used. These multipotent mesenchymal progenitor cells differentiate into osteoblasts upon treatment with the SMO agonist Purmorphamine^36^. During differentiation osteoblast specific genes such as alkaline phosphatase, which plays an essential role in bone formation, are highly expressed. Activity of alkaline phosphatase can directly be monitored by following substrate hydrolysis yielding a highly luminescent product. Inhibition of the pathway results in reduction of luminescence. Shortly, 800 cells per well were seeded in white 384 well plates (Greiner) in 25 µL medium (high glucose DMEM, 10% heat inactivated fetal calf serum, 1 mM sodium pyruvate, 6 mM L-glutamine, 100 U/mL penicillin and 0,1 mg/mL streptomycin) and allowed to grow overnight. The next day, compounds were added to a final concentration of 10 µM using the acoustic nanoliter dispenser ECHO 520 (Beckman). After one hour, 10 µL of Purmorphamine in medium were added to a final concentration of 1.5 µM using Multidrop Combi (Thermofisher Scientific); control cells were treated with DMSO. After four days, the cell culture medium was aspirated using the Elx405 cell washer (Biotek) and 25 µL of a commercial luminogenic ALK substrate (CDP-Star, Roche) were added. After one hour, luminescence was read. To identify and exclude toxic compounds that also lead to a reduction in the luminescent signal, cell viability measurements were carried out in parallel. The cell viability assay followed the same workflow as the HH assay, except that only 200 cells per well were seeded. Cell culture medium alone served as control for the cell viability assay. For the measurement of cell viability, 15 µL of CellTiterGlo reagent (Promega) which determines the cellular ATP content were added after aspiration of the medium. All data was normalized to DMSO-treated cells. Dose-response analysis was done using a three-fold dilution curve starting from 10 µM. IC_50_ values were calculated using the Quattro software suite (Quattro Research GmbH).

### GLI-dependent reporter gene assay

For evaluation of a direct effect on the Hh signalling pathway, a reporter gene assay using Shh-LIGHT2 cells was carried out.^29^. Shortly, 7500 cells per well were seeded in white 384 well plates (Greiner) in 25 µL medium (high glucose DMEM, 10% heat inactivated fetal calf serum, 1 mM sodium pyruvate, 3 mM L-glutamine, 5 mM Hepes pH 7.4, 100 U/mL penicillin and 0,1 mg/mL streptomycin) and allowed to grow overnight. The next day, compounds were added to a final concentration of 10 µM using the acoustic nanoliter dispenser ECHO 520 (Beckman). After one hour, 10 µL of Purmorphamine in medium were added to a final concentration of 3 µM using Multidrop Combi (Thermofisher Scientific); control cells were treated with DMSO. After 48 hours, 35 µL of OneGlo reagent (Promega) were added and luminescence was read. Cell viability measurements were carried out in parallel, using 750 cells/well and read-out by CellTiterGlo reagent. All data was normalized to DMSO-treated cells. Dose-response analysis was done using a three-fold dilution curve starting from 10 µM. IC_50_ values were calculated using the Quattro software suite (Quattro Research GmbH).

### Smoothened binding assay

A previously described method was modified to use the Smoothened binding assay^37^. In a 24- well plate with poly-D-lysine-coated coverslips (Neuvitro, 12 mm, #GG-12-1.5-PDL), 6 × 10^4^ HEK293T cells were seeded for 24 hours at 37 °C with 5% CO_2_. According to the manufacturer’s instructions, the cells were transfected with the SMO-expressing plasmid (pGEN_mSMO, Addgene #37673^29^) in OptiMEM medium using FuGENE® HD transfection reagent (Promega, # E2311). The plate was then kept at 37 °C in 5% CO_2_ for 48 h. The cells were then washed once with PBS, fixed with 3.7% paraformaldehyde in PBS for 10 min at room temperature, and treated for 5 min with PBS containing 10 mM glycine and 0.2% sodium azide. The fixed cells were then washed with PBS (3 times for 5 min) and treated with the compounds, Vismodegib (Selleckchem #1082) and DMSO in DMEM containing 0.5% FBS (assay medium) and 5 nM BODIPY-Cyclopamine S26 (Carbosynth Limited, FB18988) for 4 h at room temperature in dark. After that, cover slips were washed with PBS and incubated for 10 min at room temperature with 1 g/mL 4’,6 diamidino-2-phenylindole (DAPI, Sigma Aldrich, Roche, #10236276001) in PBS. Cover slips were then washed again and mounted onto glass slides using Aqua Polymount (Polysciences). Fluorescence microscopy, Zeiss Observer Z1 (Carl Zeiss, Germany) was used to acquire the images using a Plan-Apochromat 63x/1.40 Oil DIC M27 objective.

### Pan-assay interference and compound purity and

The screened compounds represent a subset of the Dark Chemical Matter that is defined as compounds that are inactive in more than 100 assays^8^ and, therefore, pan-assay interference can be excluded. All tested compounds were obtained from commercial sources.

## Acknowledgments

Research at the Max Planck Institute of Molecular Physiology was supported by the Max Planck Society. This work was co-funded by the European Union (Drug Discovery Hub Dortmund (DDHD), EFRE-0200481) and Innovative Medicines Initiative (grant agreement number 115489) resources of which are composed of financial contribution from the European Union’s Seventh Framework Programme (FP7/2007-2013) and EFPIA companies’ in-kind contribution. This work was funded from the programme “ Netzwerke 2021”, an initiative of the Ministry of Culture and Science of the State of Northrhine Westphalia. The compound management and screening center (COMAS) in Dortmund is acknowledged for performing the Cell painting assay.

## Author contributions

H.W. and S. Z. designed the research. A.P. analyzed the DCM and CPA data. J.L., S.P., S.R-A., B.S., J.W., J.B. and S.K. performed biological experiments. S.S. and A.P. performed CPA and processed the data. S.Z. analyzed data and designed experiments. A.P., S.S. and S.Z. wrote the manuscript. All authors discussed the results and commented on the manuscript.

## References

[1] Thorne, N.; Auld, D. S.; Inglese, J., Apparent Activity in High-Throughput Screening: Origins of Compound-Dependent Assay Interference, Curr Opin Chem Biol 2010, 14, 315–324.

[2] Baell, J. B.; Holloway, G. A., New Substructure Filters for Removal of Pan Assay Interference Compounds (Pains) from Screening Libraries and for Their Exclusion in Bioassays, J Med Chem 2010, 53, 2719–2740.

[3] Wetzel, S.; Bon, R. S.; Kumar, K.; Waldmann, H., Biology-Oriented Synthesis, Angew Chem Int Ed Engl 2011, 50, 10800–10826.

[4] van Hattum, H.; Waldmann, H., Biology-Oriented Synthesis: Harnessing the Power of Evolution, J Am Chem Soc 2014, 136, 11853–11859.

[5] Schreiber, S. L., Organic Chemistry: Molecular Diversity by Design, Nature 2009, 457, 153–154.

[6] Grigalunas, M.; Brakmann, S.; Waldmann, H., Chemical Evolution of Natural Product Structure, J Am Chem Soc 2022, 144, 3314–3329.

[7] Karageorgis, G.; Foley, D. J.; Laraia, L.; Waldmann, H., Principle and Design of Pseudo-Natural Products, Nat Chem 2020, 12, 227–235.

[8] Wassermann, A. M.; Lounkine, E.; Hoepfner, D.; Le Goff, G.; King, F. J.; Studer, C.; Peltier, J. M.; Grippo, M. L.; Prindle, V.; Tao, J.; Schuffenhauer, A.; Wallace, I. M.; Chen, S.; Krastel, P.; Cobos-Correa, A.; Parker, C. N.; Davies, J. W.; Glick, M., Dark Chemical Matter as a Promising Starting Point for Drug Lead Discovery, Nat Chem Biol 2015, 11, 958–966.

[9] Pope, A., Screening Heursitcs and Chemical Propery Bias; New Directions for Lead Identification and Optimization, Presented at the Society for Laboratory Automation and Screening (SLAS) Meeting, San Diego, CA, February 4−8, 2012 2012, https://www.slideshare.net/andypopeuk/screening-heuristics-popefinal.

[10] Wassermann, A. M.; Tudor, M.; Glick, M., Deorphanization Strategies for Dark Chemical Matter, Drug Discov Today Technol 2017, 23, 69–74.

[11] Gustafsdottir, S. M.; Ljosa, V.; Sokolnicki, K. L.; Wilson, J. A.; Walpita, D.; Kemp, M. M.; Seiler, K. P.; Carrel, H. A.; Golub, T. R.; Schreiber, S. L.; Clemons, P. A.; Carpenter, A. E.; Shamji, A. F., Multiplex Cytological Profiling Assay to Measure Diverse Cellular States, Plos One 2013, 8, https://doi.org/10.1371/journal.pone.0080999.

[12] Bray, M. A.; Singh, S.; Han, H.; Davis, C. T.; Borgeson, B.; Hartland, C.; Kost-Alimova, M.; Gustafsdottir, S. M.; Gibson, C. C.; Carpenter, A. E., Cell Painting, a High-Content Image-Based Assay for Morphological Profiling Using Multiplexed Fluorescent Dyes, Nat Protoc 2016, 11, 1757–1774.

[13] Carpenter, A. E.; Jones, T. R.; Lamprecht, M. R.; Clarke, C.; Kang, I. H.; Friman, O.; Guertin, D. A.; Chang, J. H.; Lindquist, R. A.; Moffat, J.; Golland, P.; Sabatini, D. M., Cellprofiler: Image Analysis Software for Identifying and Quantifying Cell Phenotypes, Genome Biol 2006, 7, R100.

[14] Christoforow, A.; Wilke, J.; Binici, A.; Pahl, A.; Ostermann, C.; Sievers, S.; Waldmann, H., Design, Synthesis, and Phenotypic Profiling of Pyrano-Furo-Pyridone Pseudo Natural Products, Angew Chem Int Ed Engl 2019, 58, 14715–14723.

[15] Pahl, A.; Schölermann, B.; Rusch, M.; Dow, M.; Hedberg, C.; Nelson, A.; Sievers, S.; Waldmann, H.; Ziegler, S., Morphological Subprofile Analysis for Bioactivity Annotation of Small Molecules, bioRxiv 2022, 2022.2008.2015.503944.

[16] Schneidewind, T.; Brause, A.; Scholermann, B.; Sievers, S.; Pahl, A.; Sankar, M. G.; Winzker, M.; Janning, P.; Kumar, K.; Ziegler, S.; Waldmann, H., Combined Morphological and Proteome Profiling Reveals Target-Independent Impairment of Cholesterol Homeostasis, Cell Chem Biol 2021, 28, 1780–1794 e1785.

[17] Nadanaciva, S.; Lu, S. Y.; Gebhard, D. F.; Jessen, B. A.; Pennie, W. D.; Will, Y., A High Content Screening Assay for Identifying Lysosomotropic Compounds, Toxicol in Vitro 2011, 25, 715–723.

[18] Keiser, M. J.; Roth, B. L.; Armbruster, B. N.; Ernsberger, P.; Irwin, J. J.; Shoichet, B. K., Relating Protein Pharmacology by Ligand Chemistry, Nat Biotechnol 2007, 25, 197–206.

[19] Comess, K. M.; McLoughlin, S. M.; Oyer, J. A.; Richardson, P. L.; Stockmann, H.; Vasudevan, A.; Warder, S. E., Emerging Approaches for the Identification of Protein Targets of Small Molecules - a Practitioners’ Perspective, J Med Chem 2018, 61, 8504–8535.

[20] Altmann, K. H.; Wartmann, M.; O’Reilly, T., Epothilones and Related Structures--a New Class of Microtubule Inhibitors with Potent in Vivo Antitumor Activity, Biochim Biophys Acta 2000, 1470, M79–91.

[21] Lacey, E.; Watson, T. R., Structure-Activity-Relationships of Benzimidazole Carbamates as Inhibitors of Mammalian Tubulin, Invitro, Biochem Pharmacol 1985, 34, 1073–1077.

[22] Shipman, C., Jr.; Smith, S. H.; Carlson, R. H.; Drach, J. C., Antiviral Activity of Arabinosyladenine and Arabinosylhypoxanthine in Herpes Simplex Virus-Infected Kb Cells: Selective Inhibition of Viral Deoxyribonucleic Acid Synthesis in Synchronized Suspension Cultures, Antimicrob Agents Chemother 1976, 9, 120–127.

[23] Fredholm, B. B., Adenosine, an Endogenous Distress Signal, Modulates Tissue Damage and Repair, Cell Death Differ 2007, 14, 1315–1323.

[24] Schneidewind, T.; Brause, A.; Pahl, A.; Burhop, A.; Mejuch, T.; Sievers, S.; Waldmann, H.; Ziegler, S., Morphological Profiling Identifies a Common Mode of Action for Small Molecules with Different Targets, Chembiochem 2020, 21, 3197–3207.

[25] Bramson, H. N.; Corona, J.; Davis, S. T.; Dickerson, S. H.; Edelstein, M.; Frye, S. V.; Gampe, R. T., Jr.; Harris, P. A.; Hassell, A.; Holmes, W. D.; Hunter, R. N.; Lackey, K. E.; Lovejoy, B.; Luzzio, M. J.; Montana, V.; Rocque, W. J.; Rusnak, D.; Shewchuk, L.; Veal, J. M.; Walker, D. H.; Kuyper, L. F., Oxindole-Based Inhibitors of Cyclin-Dependent Kinase 2 (Cdk2): Design, Synthesis, Enzymatic Activities, and X-Ray Crystallographic Analysis, J Med Chem 2001, 44, 4339–4358.

[26] Schoelermann, B.; Bonowski, J.; Grigalunas, M.; Burhop, A.; Xie, Y.; Hoock, J. G. F.; Liu, J.; Dow, M.; Nelson, A.; Nowak, C.; Pahl, A.; Sievers, S.; Ziegler, S., Identification of Dihydroorotate Dehydrogenase Inhibitors Using the Cell Painting Assay, Chembiochem 2022, 23, e202200475.

[27] Vincent, F.; Loria, P.; Pregel, M.; Stanton, R.; Kitching, L.; Nocka, K.; Doyonnas, R.; Steppan, C.; Gilbert, A.; Schroeter, T.; Peakman, M. C., Developing Predictive Assays: The Phenotypic Screening “Rule of 3”, Sci Transl Med 2015, 7, 293ps215.

[28] Flegel, J.; Shaaban, S.; Jia, Z. J.; Schulte, B.; Lian, Y.; Krzyzanowski, A.; Metz, M.; Schneidewind, T.; Wesseler, F.; Flegel, A.; Reich, A.; Brause, A.; Xue, G.; Zhang, M.; Dotsch, L.; Stender, I. D.; Hoffmann, J. E.; Scheel, R.; Janning, P.; Rastinejad, F.; Schade, D.; Strohmann, C.; Antonchick, A. P.; Sievers, S.; Moura-Alves, P.; Ziegler, S.; Waldmann, H., The Highly Potent Ahr Agonist Picoberin Modulates Hh-Dependent Osteoblast Differentiation, J Med Chem 2022, 65, 16268–16289.

[29] Taipale, J.; Chen, J. K.; Cooper, M. K.; Wang, B.; Mann, R. K.; Milenkovic, L.; Scott, M. P.; Beachy, P. A., Effects of Oncogenic Mutations in Smoothened and Patched Can Be Reversed by Cyclopamine, Nature 2000, 406, 1005–1009.

[30] Wu, F.; Zhang, Y.; Sun, B.; McMahon, A. P.; Wang, Y., Hedgehog Signaling: From Basic Biology to Cancer Therapy, Cell Chem Biol 2017, 24, 252–280.

[31] Scholermann, B.; Bonowski, J.; Grigalunas, M.; Burhop, A.; Xie, Y.; Hoock, J. G. F.; Liu, J.; Dow, M.; Nelson, A.; Nowak, C.; Pahl, A.; Sievers, S.; Ziegler, S., Identification of Dihydroorotate Dehydrogenase Inhibitors Using the Cell Painting Assay, Chembiochem 2022, 23, e202200475.

[32] Bray, M. A.; Singh, S.; Han, H.; Davis, C. T.; Borgeson, B.; Hartland, C.; Kost-Alimova, M.; Gustafsdottir, S. M.; Gibson, C. C.; Carpenter, A. E., Cell Painting, a High-Content Image-Based Assay for Morphological Profiling Using Multiplexed Fluorescent Dyes, Nat Protoc 2016, 11, 1757–1774.

[33] Woehrmann, M. H.; Bray, W. M.; Durbin, J. K.; Nisam, S. C.; Michael, A. K.; Glassey, E.; Stuart, J. M.; Lokey, R. S., Large-Scale Cytological Profiling for Functional Analysis of Bioactive Compounds, Mol Biosyst 2013, 9, 2604–2617.

[34] Grigalunas, M.; Burhop, A.; Zinken, S.; Pahl, A.; Gally, J. M.; Wild, N.; Mantel, Y.; Sievers, S.; Foley, D. J.; Scheel, R.; Strohmann, C.; Antonchick, A. P.; Waldmann, H., Natural Product Fragment Combination to Performance-Diverse Pseudo-Natural Products, Nat Commun 2021, 12, 1883.

[35] Akbarzadeh, M.; Deipenwisch, I.; Schoelermann, B.; Pahl, A.; Sievers, S.; Ziegler, S.; Waldmann, H., Morphological Profiling by Means of the Cell Painting Assay Enables Identification of Tubulin-Targeting Compounds, Cell Chem Biol 2022, 29, 1053–1064 e1053.

[36] Wu, X.; Ding, S.; Ding, Q.; Gray, N. S.; Schultz, P. G., A Small Molecule with Osteogenesis-Inducing Activity in Multipotent Mesenchymal Progenitor Cells, J Am Chem Soc 2002, 124, 14520–14521.

[37] Sinha, S.; Chen, J. K., Purmorphamine Activates the Hedgehog Pathway by Targeting Smoothened, Nat Chem Biol 2006, 2, 29–30.

