## Supporting Information for "Illuminating Dark Chemical Matter using the Cell Painting Assay"

#### Table of contents

Supporting Figures ........ S2

Supporting Tables ........ S9

References ........ S13

### Supporting Figures

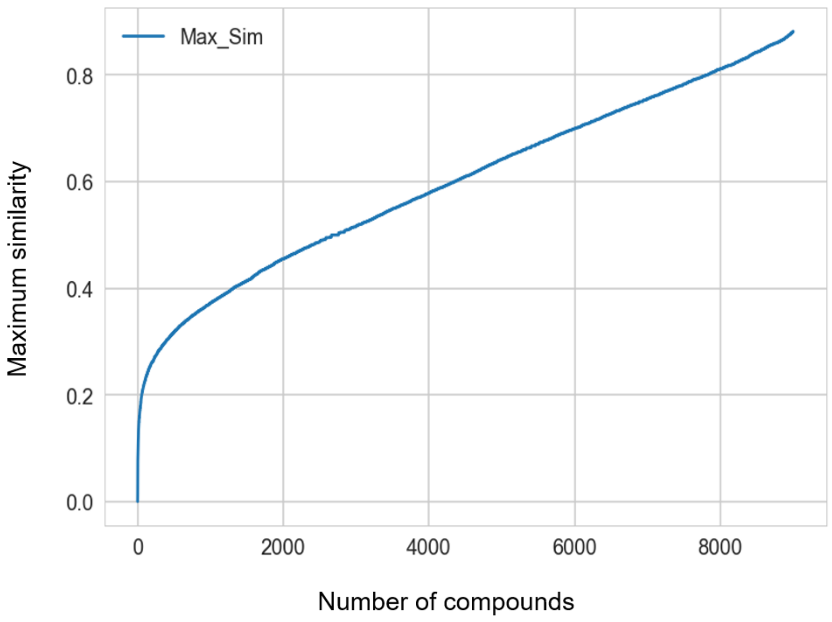


**Figure S1 (related to Figure 1):** Increasing maximum chemical similarity of compounds added to the set during the diversity selection. The similarity was determined by Tanimoto similarity of the RDKit Morgan fingerprints (radius 2).

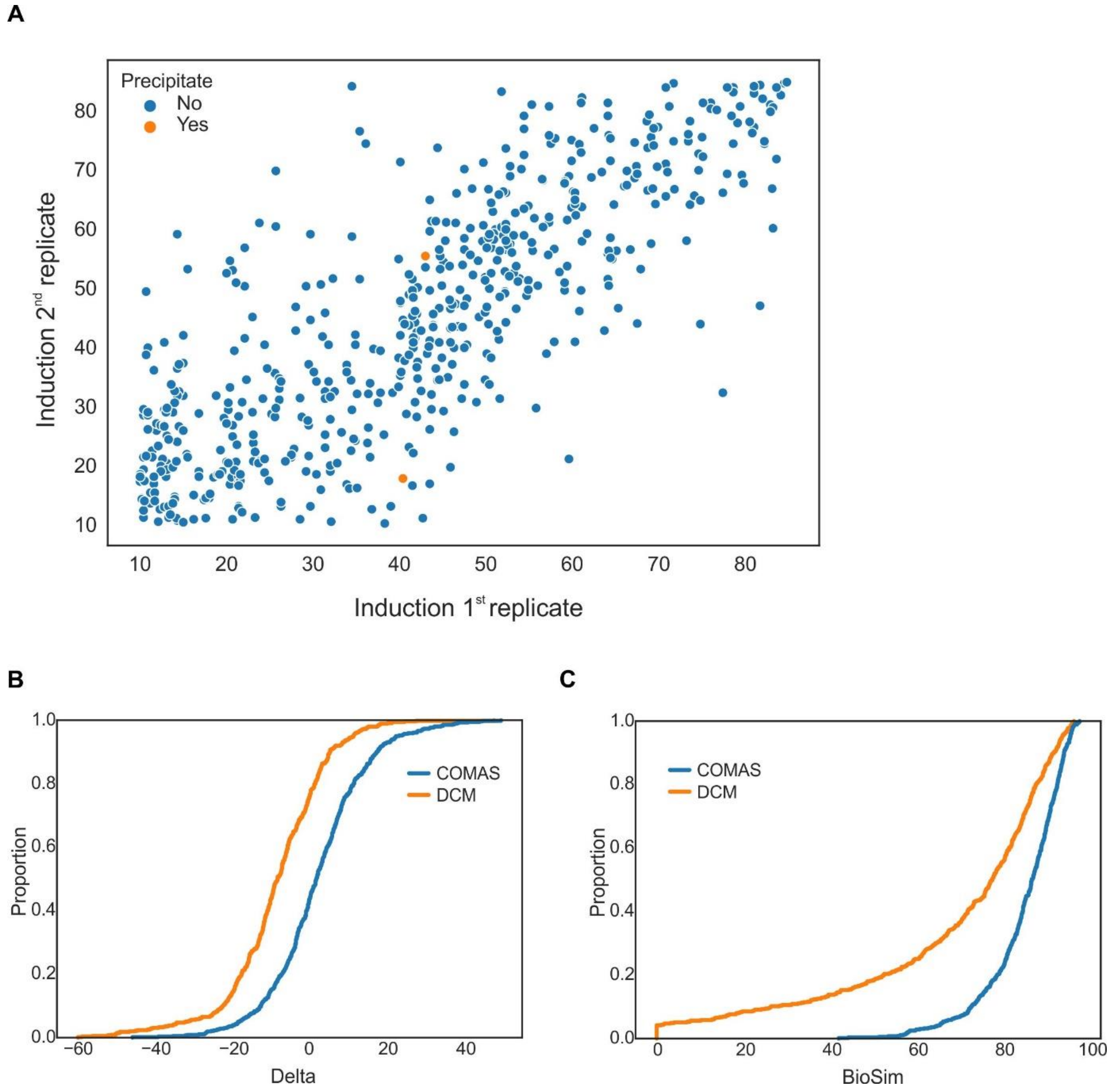


**Figure S2 (related to Figure 1): Reproducibility for DCM in the Cell painting.** (A) Reproducibility of the induced changes as determined by the induction value for two replicates for 562 CPA-active internal compounds. Correlation between the replicates: $r^2$=0.617. (B) Empirical cumulative distribution function for differences in induction (2nd replicate – 1st replicate) of 562 internal compounds and the 550 DCM compounds. (C) Empirical cumulative distribution functions for the profile similarity (biosimilarity, BioSim) between the two biological replicates for the 562 internal and 550 DCM compounds.

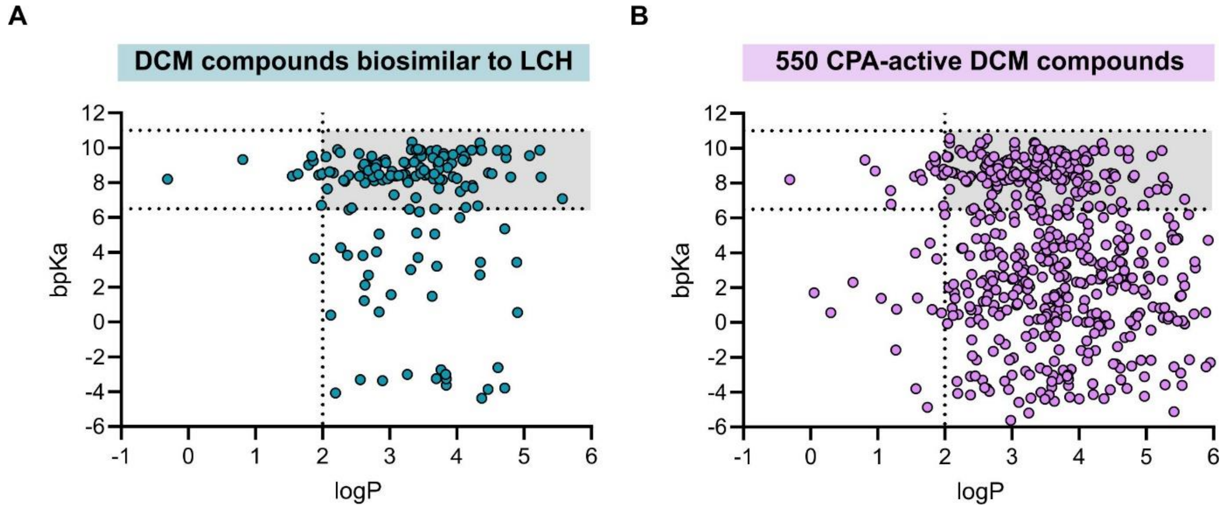


**Figure S3 (related to Figure 1).** Calculated logP and bpKa values CPA-active DCM compounds that show subprofile biosimilarity ≥ 80% to the lysosomotropism/cholesterol homeostasis cluster (LCH) (A). (B) logP and bpKa values for all 550 active DCM compounds The grey region corresponds to logP > 2 and bpKa between 6.5 and 11 and to properties of lysosomotropic compounds [1].

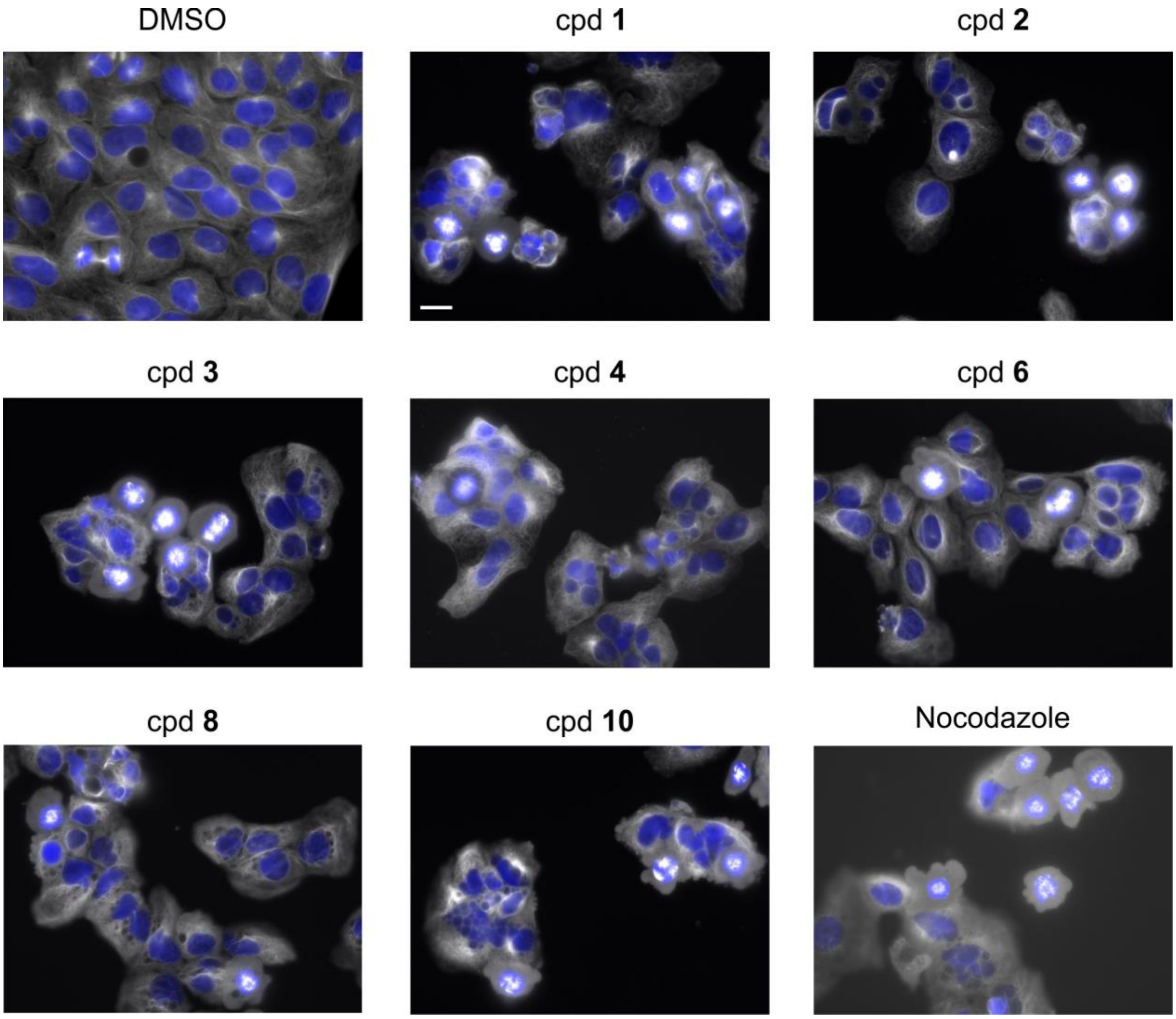


**Figure S4. Influence of compounds on the microtubule cytoskeleton.** U2OS cells were treated with 30 µM of the compound or DMSO and Nocodazole (0.1 µM) as controls for 24 h prior to staining with anti-tubulin antibody (white) or DAPI (blue) to visualize the DNA. Scale bar: 20 µm.

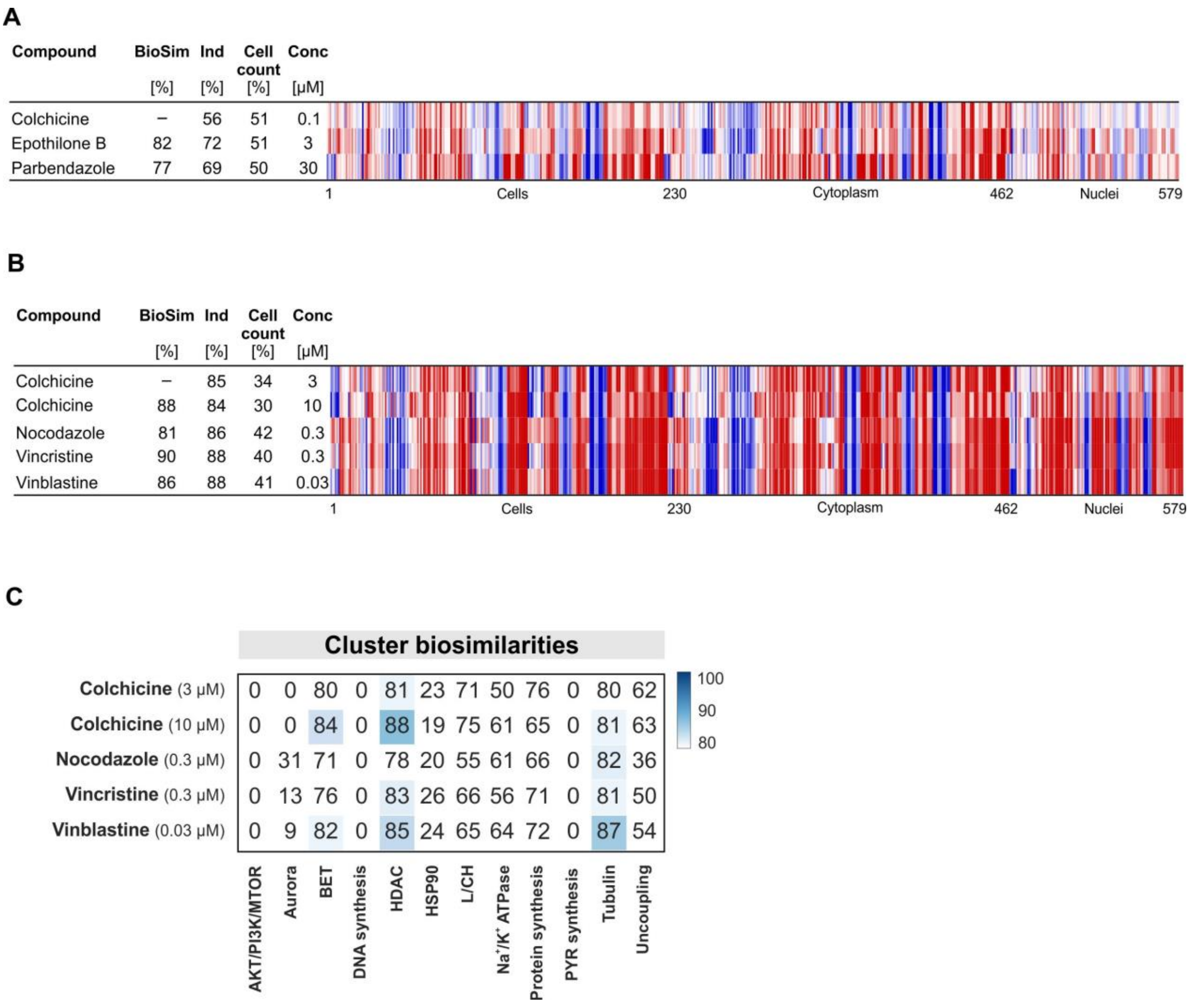


**Figure S5. CPA analysis for Colchicine.** (A and B) Biosimilarity of Colchicine to Epothilone and Parbendazole (A) or to Nocodazole, Vincristine and Vinblastine at concentration with cell count < 50%. (B). The top line profile is set as a reference profile (100 % biological similarity, BioSim) to which the following profiles are compared. Blue color: decreased feature, red color: increased feature. BioSim: biosimilarity, Ind: induction, Conc: concentration. (C) Cluster biosimilarity heatmap for Colchicine, Nocodazole, Vincristine and Vinblastine at a concentration with cell count < 50%.

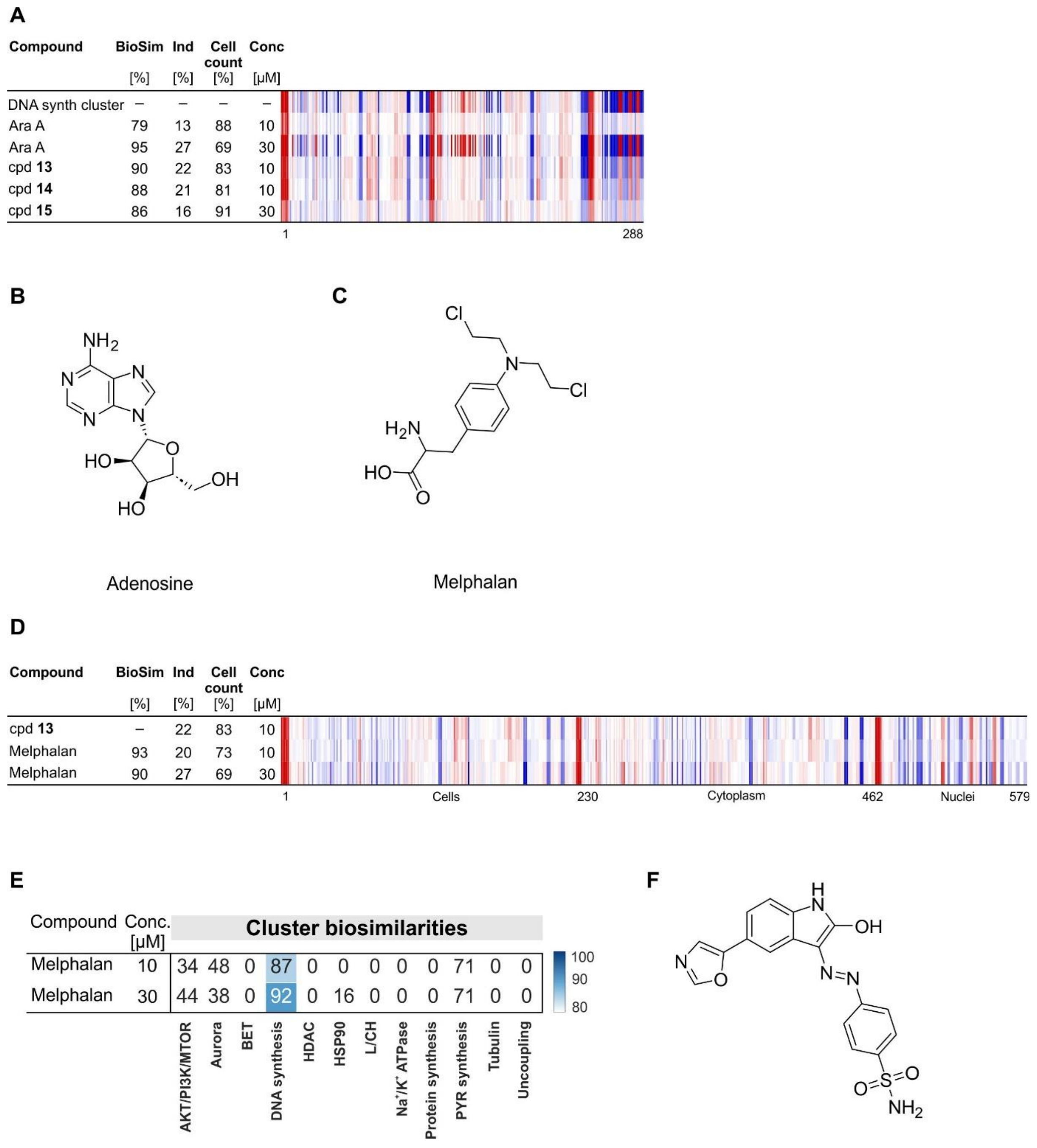


**Figure S6 (related to Figure 4): DCM compound with similarity to the DNA synthesis cluster.** (A) Biosimilarity to the DNA synthesis (synth) cluster profile. The top line profile is set as a reference profile (100 % biological similarity, BioSim) to which the following profiles are compared. Blue color: decreased feature, red color: increased feature. BioSim: biosimilarity, Ind: induction, Conc: concentration. (B) Structure of adenosine. (C) Structure of Melphalan. (D) Profile similarity of compound (cpd) **13** to Melphalan. The top line profile is set as a reference profile (100 % biological similarity, BioSim) to which the following profiles are compared. Blue color: decreased feature, red color: increased feature. BioSim: biosimilarity,

Ind: induction, Conc: concentration. (E) Cluster biosimilarity for Melphalan. PYR: pyrimidine. (F) Structure of the oxindole-based DCK inhibitor from Figure 4D.

### Supporting Tables

**Table S1:** See separate XLS file

**Table S2 Related to Figure 5. DCM compounds that suppress Hedgehog-induced osteogenesis.** $IC_{50}$ values ± SD (n=3) for inhibition of Hedgehog-induced osteogenesis in C3H10T1/2 cells or GLI-dependent reporter gene assay in Shh-LIGHT2 cells (GLI RGA).

| Cpd | Structure | Osteogenesis inhibition $IC_{50}$ ± SD | GLI RGA inhibition $IC_{50}$ ± SD |
|---|---|---|---|
| **17** | 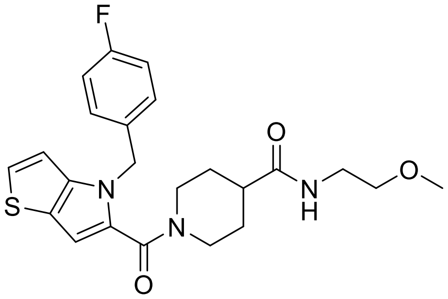 | 0.09 ± 0 | 0.47 ± 0.06 |
| **18** | 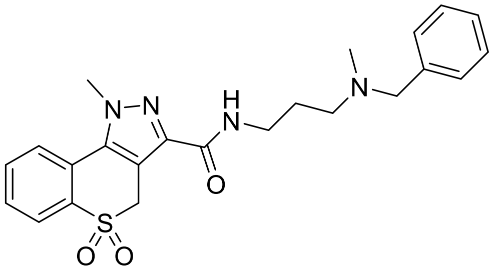 | 0.31 ± 0.02 | 2.17 ± 0.2 |
| **19** | 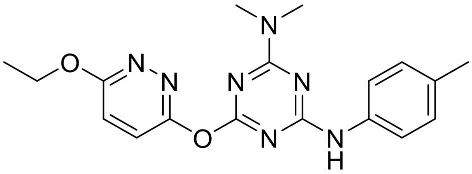 | 0.32 ± 0.2 | |
| **20** | 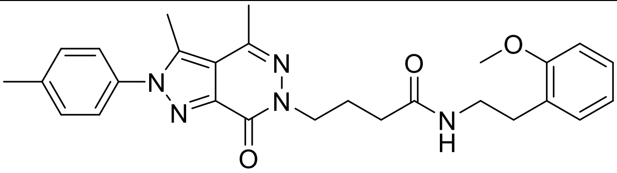 | 0.54 ± 0.02 | 1.40 ± 0.1 |
| **21** | 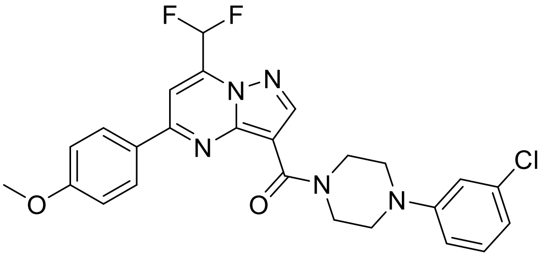 | 0.92 ± 0.29 | 1.51 ± 0.2 |

| Cpd | Structure | Osteogenesis inhibition $IC_{50} \pm SD$ | GLI RGA inhibition $IC_{50} \pm SD$ |
|---|---|---|---|
| **22** | 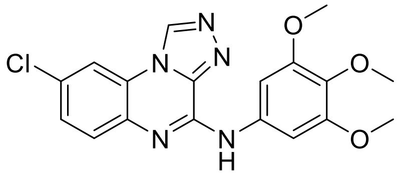 | 1.16 ± 0.1 | 1.02 ± 0.2 |
| **23** | 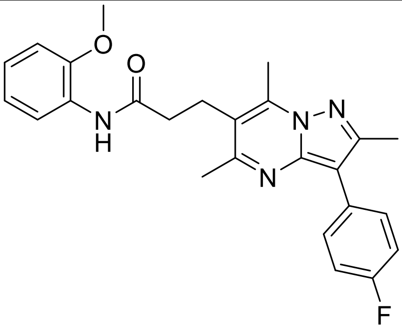 | 1.28 ± 0.4 | 6.90 ± 0.8 |
| **24** | 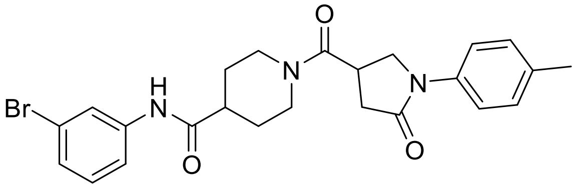 | 1.38 ± 0.3 | 7.99 ± 0.0 |
| **25** | 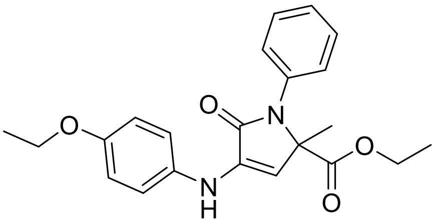 | 1.46 ± 0.4 | |
| **26** | 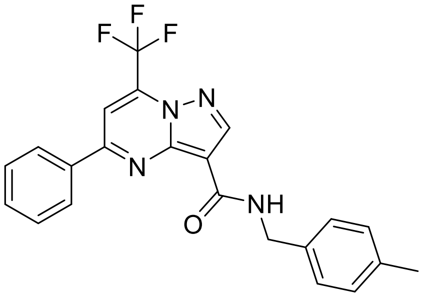 | 1.57 ± 0.3 | |
| **27** | 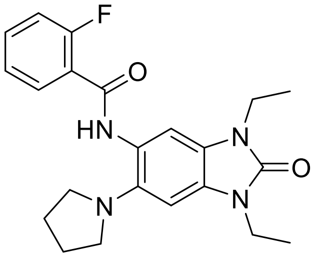 | 1.66 ± 0.6 | |
| **28** | 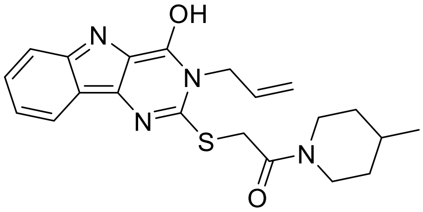 | 2.02 ± 0.3 | |

| Cpd | Structure | Osteogenesis inhibition $IC_{50} \pm SD$ | GLI RGA inhibition $IC_{50} \pm SD$ |
|---|---|---|---|
| **29** | 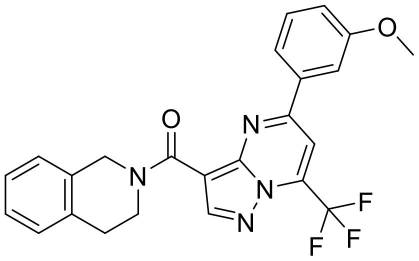 | 2.13 ± 0.3 | |
| **30** | 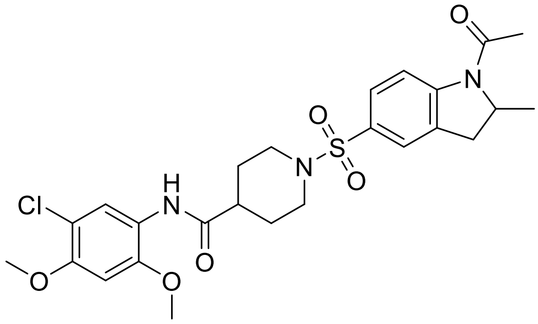 | 2.35 ± 0.2 | |
| **31** | 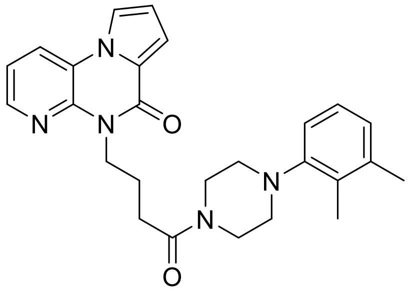 | 3.36 ± 0.2 | |
| **32** | 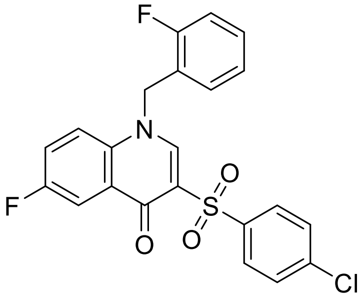 | 3.89 ± 0.6 | |
| **33** | 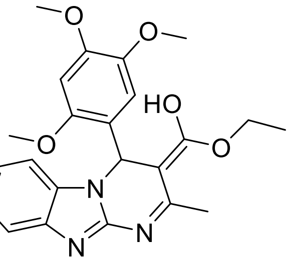 | 4.01 ± 0.6 | |
| **34** | 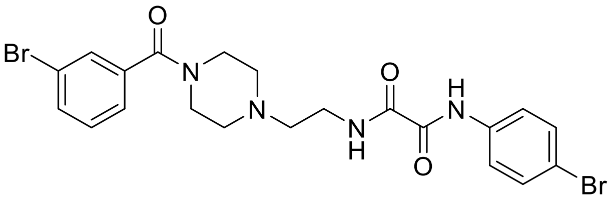 | 4.25 ± 0.4 | |

| Cpd | Structure | Osteogenesis inhibition $IC_{50}$ ± SD | GLI RGA inhibition $IC_{50}$ ± SD |
|---|---|---|---|
| **3935** | 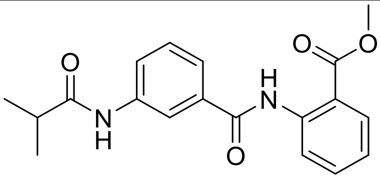 | 4.36 ± 1.3 | |
| **36** | 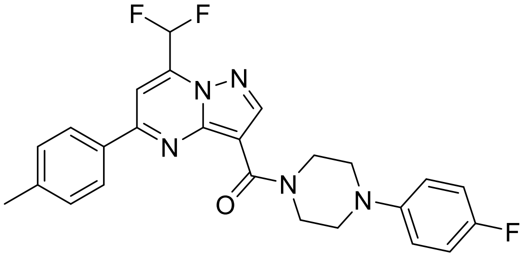 | 4.55 ± 1.4 | |
| **37** | 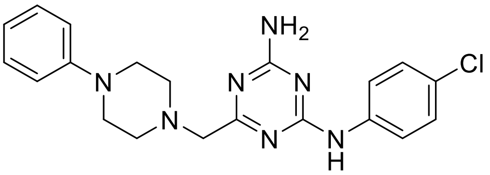 | 5.95 ± 0.6 | |
| **38** | 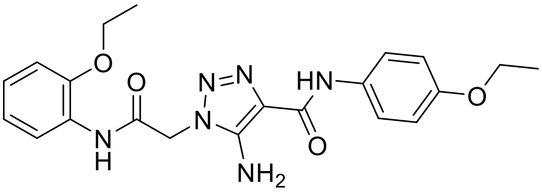 | 6.87 ± 1.3 | |
| **39** | 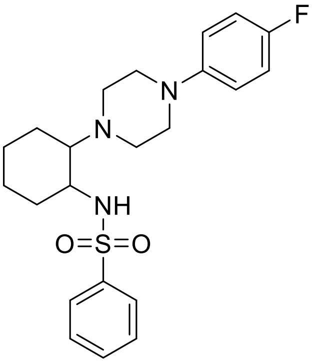 | 7.00 ± 0.6 | |
| **40** | 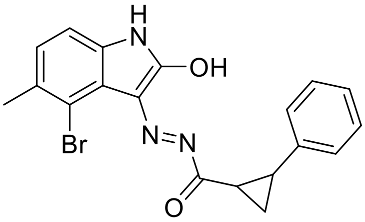 | 7.46 ± 0.0 | |

### References

[1] Nadanaciva, S.;Lu, S. Y.;Gebhard, D. F.;Jessen, B. A.;Pennie, W. D.; Will, Y., A High Content Screening Assay for Identifying Lysosomotropic Compounds, *Toxicol in Vitro* **2011**, *25*, 715-723.
